# The herpesvirus UL49.5 protein hijacks a cellular C-degron pathway to drive TAP transporter degradation

**DOI:** 10.1101/2023.09.27.559663

**Authors:** Magda Wąchalska, Celeste Riepe, Magdalena J. Ślusarz, Małgorzata Graul, Lukasz S. Borowski, Wenjie Qiao, Michalina Foltynska, Jan E. Carette, Krystyna Bieńkowska-Szewczyk, Roman J. Szczesny, Ron R. Kopito, Andrea D. Lipińska

## Abstract

The transporter associated with antigen processing (TAP) is a key player in the MHC class I-restricted antigen presentation and an attractive target for immune evasion by viruses. Bovine herpesvirus 1 (BoHV-1) impairs TAP-dependent antigenic peptide transport through a two-pronged mechanism in which binding of the UL49.5 gene product to TAP both inhibits peptide transport and promotes its proteasomal degradation. How UL49.5 promotes TAP degradation is unknown. Here, we use high-content siRNA and genome-wide CRISPR-Cas9 screening to identify CLR2^KLHDC3^ as the E3 ligase responsible for UL49.5-triggered TAP disposal in human cells. We propose that the C-terminus of UL49.5 mimics a C-end rule degron that recruits the E3 to TAP and engages the CRL2 E3 in ER-associated degradation.

**SIGNIFICANCE:** Herpesviruses are masters of immune evasion. Most often, they hijack host cellular pathways to modulate the antiviral immune response. Varicellovirus UL49.5 orthologs have evolved as inhibitors of the transporter associated with antigen processing (TAP) and, this way, major modulators of the MHC class I-restricted antigen presentation. This study identifies the long-sought molecular mechanism exploited by bovine herpesvirus 1-encoded UL49.5 to trigger proteasomal degradation of TAP. Our findings demonstrate that the viral protein hijacks host cell CRL2-ubiquitin conjugation and ER-associated degradation pathways to promote TAP degradation. These findings advance the understanding of how herpesviruses can manipulate the cellular machinery.

## INTRODUCTION

Herpesviruses have achieved success in persistent and recurrent infections due to their remarkable immunomodulatory properties. These viruses have developed distinct mechanisms to evade both innate and adaptive immune responses as they co-evolved with their hosts. Downregulation of major histocompatibility (MHC) class I-dependent antigen presentation is one of the most efficient strategies to escape cytotoxic CD8^+^ T lymphocytes (CTLs). A key player in this pathway - the transporter associated with antigen processing (TAP) - is an attractive target for inhibition by many herpesviruses (1). The role of TAP is to transport antigenic peptides from the cytoplasm to the lumen of the endoplasmic reticulum (ER), where they are loaded on MHC I and subsequently displayed on the cell surface to CTLs (2,3). TAP is a heterodimer belonging to the ATP-binding cassette family transporters. It consists of two subunits, TAP1 and TAP2. The core of each subunit is formed by an N-terminal transmembrane domain (TMD) responsible for peptide recognition and binding and a highly conserved C-terminal nucleotide-binding domain (NBD), which can bind and hydrolyze ATP (4–6). The mechanism of TAP inactivation by bovine herpesvirus 1 (BoHV-1) is unique among viruses, as direct binding of the virally-encoded BoHV-1 UL49.5 protein to TAP both restrains the conformational cycle required for peptide transport and directs TAP for proteasomal degradation (7).

UL49.5 is a ∼10 kDa type I transmembrane protein comprised of a predicted 32 amino acid (aa) non-glycosylated extracellular domain and a 17 aa cytosolic tail that is essential to promote TAP degradation but dispensable for inhibition of TAP-mediated peptide transport (**Figure 1A and 1B**) (7–9). UL49.5 binds directly to the TAP1/TAP2 heterodimer and triggers the degradation of both TAP subunits in a process requiring valosin-containing protein (VCP)/p97 (10), an AAA^+^ ATPase that facilitates the dislocation of membrane-integrated substrates of ER-associated degradation (ERAD) (11–13).

**Figure 1.**
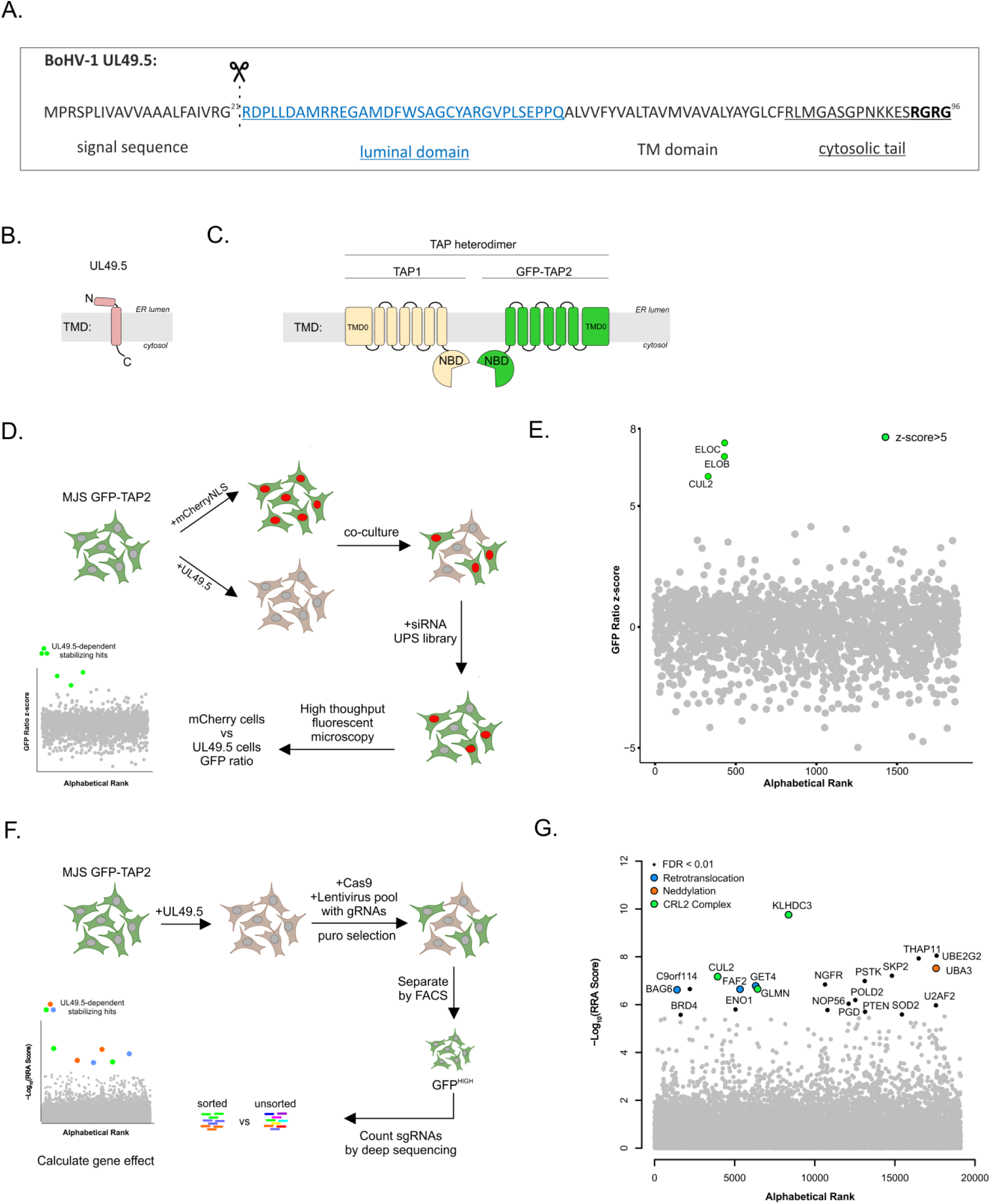
siRNA and CRISPR-Cas9 screens identify cellular factors utilized by UL49.5 for TAP degradation. (A) UL49.5 protein sequence with the predicted domains(9) (the signal peptidase cleavage site is denoted). (B) Schematic representation of UL49.5 topology in the ER membrane and domain organization. TMD: transmembrane domain. (C) Schematic of the fluorescent TAP model. NBD: nucleotide-binding domain;TMD: transmembrane domain;TMD0: additional transmembrane domain (D) siRNA screen workflow. (E) Significance of genes in siRNA screen based on the +/-UL49.5 GFP ratio z-score. The complete list of ranked genes is included in Table S1. (F) Genome-wide CRISPR-Cas9 screen workflow. (G) Robust rank aggregation (RRA) scores of genes observed in the CRISPR-Cas9 screen. Black: significant genes with FDR < 0.05. Labeled genes: significant genes with FDR < 0.01. Hits were colored by pathways. The complete list of ranked genes can be found in Table S2.

In principle, UL49.5 could promote TAP degradation by acting as an E3 ubiquitin ligase to directly transfer ubiquitin chains to TAP subunits. Such a strategy is employed by the murine gammaherpesvirus-68 mK3 protein and its ortholog from the rodent herpesvirus Peru (14, 15). Alternatively, UL49.5 could function as an adaptor that hijacks the host ubiquitin-proteasome system (UPS) to induce TAP ubiquitylation and subsequent proteasomal degradation. To test these hypotheses, we exploited an unbiased functional genomic approach to identify host factors that are required for UL49.5-mediated TAP degradation. These experiments support the latter hypothesis, identifying the cullin 2 ubiquitin ligase and its substrate recognition receptor, Kelch domain-containing protein 3/KLHDC3 (CRL2^KLHDC3^), as responsible for UL49.5-triggered TAP degradation via a C-degron pathway.

## RESULTS

### Functional genetic screens identify the E3 ubiquitin ligase responsible for UL49.5-induced TAP degradation

We used complementary functional genomic approaches to identify host factors that underlie UL49.5-triggered TAP degradation. We used a previously described (10) MelJuso (MJS) cell reporter line in which the endogenous *TAP2* gene is replaced with *GFP-TAP2* (**Figure 1C**). MJS cells are commonly used to investigate immunomodulatory properties of herpesviruses (16, 17) and are permissive to bovine herpesvirus (18). Control experiments confirmed that this MJS GFP-TAP2 reporter line faithfully recapitulates UL49.5-induced, UPS-dependent TAP degradation (**Figure S1A**).

We used a high-content microscopy-based siRNA screen of 1,998 UPS-related genes (**Figure 1D**) in our MJS GFP-TAP2 reporter cells. To find genes that stabilized GFP-TAP2 specifically in response to UL49.5 expression, MJS GFP-TAP2 cells transduced with either UL49.5 or mCherry-NLS were co-cultured at a 1:1 ratio and transfected with the siRNA library in an arrayed format such that each well received a pool of 4 gene-specific siRNAs. Automated microscopic image analysis was used to quantify GFP and mCherry fluorescence intensities in single cells and to exclude cells with aberrant morphologies. Comparison of GFP signals between mCherry +/-cells allowed us to identify genes whose depletion resulted in statistically significant increased GFP-TAP2 fluorescence specifically in the presence of UL49.5. We found that cullin 2 (*CUL2*) and its cognate substrate adaptors, elongins B and C (*ELOB, ELOC*), were the top hits, standing apart from the other candidates, with a z-score > 5 (**Figure 1E**). These data suggest that UL49.5 promotes TAP degradation by hijacking an endogenous cullin-RING E3 ligase (CRL2).

CRLs are modular multi-subunit E3s that are composed of a cullin scaffold, an E2-binding RING domain-containing protein (RBX1 or RBX2, depending on the CRL), substrate adaptors, and a substrate recognition receptor (SRR), which dictates the substrate selectivity of the CRL (19). While the siRNA screen identified CUL2 and its cognate elongin B/C adaptors, it failed to identify the SRR, so we expanded the search for genes involved in UL49.5 mediated TAP2 degradation by conducting a genome-wide CRISPR/Cas9 knockout screen (**Figure 1F**). We transduced MJS GFP-TAP2 cells stably expressing UL49.5 with Cas9 nuclease and introduced the human Brunello CRISPR pooled lentiviral library, which consists of 77,441 sgRNAs, with an average of 4 sgRNAs per gene (20). We used fluorescence-activated cell sorting (FACS) to collect the brightest 5% of GFP-TAP2-expressing cells, and sgRNAs from the sorted and unsorted populations were sequenced and enrichment scores for each gene were calculated using MAGeCK (**Figure 1G**). Consistent with the results of the siRNA screen, we observed strong enrichment of sgRNAs targeting *CUL2*. Importantly, the CRISPR screen added *KLHDC3* - a known cullin 2 SSR to the list of UL49.5/TAP-modulated genes, in addition to the cullin regulatory factor glomulin (GLMN) (21), and the NEDD8-activating enzyme E1 (UBA3) (22) (**Figure 1G**). Together, the two complementary screening approaches strongly implicate CRL2^KLHDC3^ in mediating UL49.5-dependent TAP degradation.

The CRISPR screen also identified FAF2, an integral membrane protein that recruits p97/VCP to the ER membrane via its cytosolic UBX domain to dislocate membrane-integrated ubiquitylated clients from the ER (11), as well as BAG6 and GET4, which function together to chaperone hydrophobic and amphipathic proteins to the proteasome downstream of FAF2 and p97/VCP (23).

### UL49.5 hijacks CRL2^KLHDC3^ to target TAP for ERAD

To validate the results of our screens, we transfected MJS GFP-TAP2 cells with siRNAs targeting selected genes. Knockdown of *CUL2* and *KLHDC3* resulted in significantly increased fluorescence of the GFP-TAP2 reporter (**Figure 2A**) as well as increased steady-state level of TAP1 and GFP-TAP2 protein only in the presence of UL49.5 (**Figure 2B**). By contrast, knockdown of the gene encoding cullin 5 (*CUL5*), which shares a similar elongin BC box with cullin 2 (24), failed to stabilize TAP (**Figure 2A, 2B** and **S2A**). We were unable to detect endogenous KLHDC3 in MJS cells with available antibodies, likely owing to its low expression (25). Therefore, we confirmed the efficacy of *KLHDC3* siRNA by transient co-transfection of N-terminally myc-tagged *KLHDC3* (^myc^KLHDC3) and siRNA targeting the *KLHDC3* gene (**Figure S2B** and **S2C**). We observed that, in the presence of UL49.5, *CUL2* and *KLHDC3* siRNA stabilized endogenous TAP1 and TAP2 in wildtype MJS cells to a similar extent to that observed in our reporter cell lines (**Figure 2C**). Moreover, *KLHDC3* depletion increased steady-state TAP1 level to that observed in the absence of UL49.5 (compare lanes 1, 5, and 7 in **Figure 2C**). Thus, the effects of depleting *CUL2* and *KLHDC3* on UL49.5-mediated TAP turnover is not an artifact of the use of GFP-tagged reporters.

**Figure 2.**
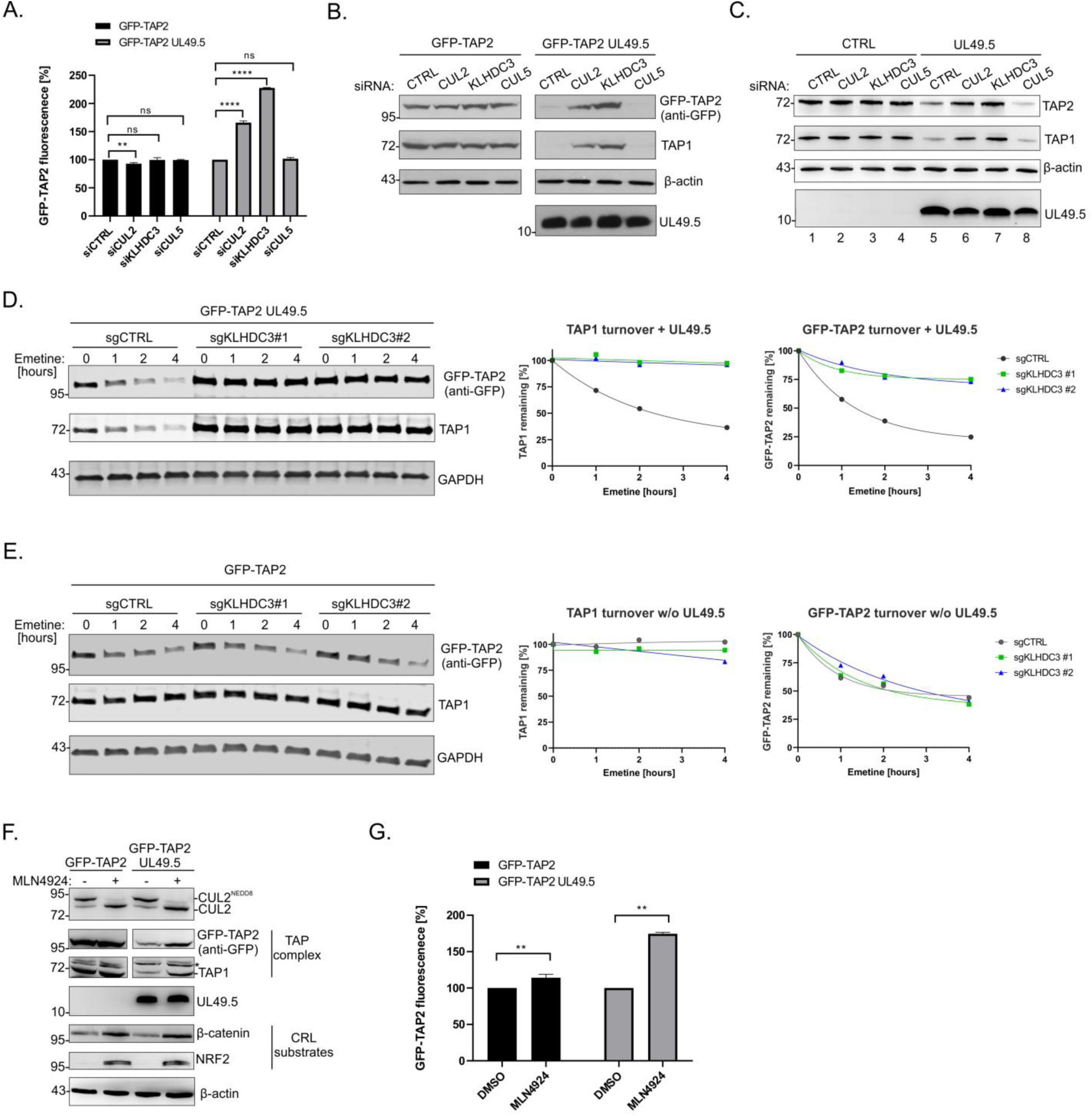
CRL2^KLHDC3^ complex is required for UL49.5-mediated TAP degradation. (A) MJS GFP-TAP2 cells stably expressing UL49.5 were transfected with the indicated siRNAs and analyzed by flow cytometry. The mean GFP fluorescence intensity from three independent measurements is represented as a percentage of the control cell (siCRTL) mean fluorescence intensity with standard deviations. The statistical significance was assessed by one-way ANOVA; **p< 0.01 ****p<0.0001. See **Figure S2A** for the efficiency of cullin depletion. (B-C) MJS GFP-TAP2 (B) and MJS (C) cells stably expressing UL49.5 were transfected with the indicated siRNAs. Steady-state levels of TAP1 and TAP2 were analyzed by immunoblotting with the indicated antibodies. See also Figure S2. (D-E) MJS GFP-TAP2 stably expressing UL49.5 (D) and MJS GFP-TAP2 (E) cells were transduced with sgRNAs targeting *KLHDC3*. The turnover of TAP1 and GFP-TAP2 was assessed by immunoblotting in the lysates of these cells collected at the indicated time points after their treatment with the protein synthesis inhibitor emetine. Quantification of TAP subunits was normalized to GAPDH. (F-G) MJS GFP-TAP2 cells stably expressing UL49.5 were treated with the neddylation inhibitor MLN4924 or DMSO as a control. The steady-state levels of TAP subunits were analyzed by immunoblotting with the indicated antibodies (F). CUL2^NEDD8^ denotes CUL2 activated by NEDD8 modification. The stability of the fluorescent reporter was analyzed by flow cytometry (G). The mean GFP fluorescence intensity from three independent measurements is represented as the percentage of the control cells, with standard deviations. The statistical significance was assessed by one-way ANOVA; **p< 0.01.

To confirm the role of KLHDC3 in UL49.5-mediated TAP degradation, we used translation shutoff assays to assess TAP subunit turnover in MJS GFP-TAP2 cells transduced with lentiviruses expressing *KLHDC3*-targeting sgRNAs (*KLHDC3* KO). In the absence of UL49.5, TAP is a stable dimer with a half-life exceeding 8 hours (7). However, in the presence of UL49.5, TAP1 and GFP-TAP2 subunits were degraded with half-lives of approximately 2 and 1.5 hours, respectively (**Figure 2D**). In the presence of UL49.5, *KLHDC3* KO strongly stabilized both TAP1 and GFP-TAP2 (**Figure 2D**). The small fraction of GFP-TAP2 that decays in *KLHDC3* KO cells most likely reflects the degradation of unassembled GFP-TAP2 which, in these transgenic cells, is expressed in excess over endogenous TAP1. Indeed, previous studies have reported the intrinsic instability of unassembled TAP2 subunits when expressed in stoichiometric excess over TAP1 (26, 27). By contrast, in the absence of UL49.5, the turnover of both TAP1 and GFP-TAP2 was unaffected by *KLHDC3* disruption (**Figure 2E**), strongly suggesting that CRL2^KLHDC3^ targets TAP subunit degradation exclusively in the presence of UL49.5 and suggesting that UL49.5 promotes TAP2 degradation by a different mechanism than the one used to destroy excess TAP subunits. The stabilization of both TAP1 and GFP-TAP2 observed upon either transient siRNA-mediated knockdown (**Figures 2A-C**) or stable gene disruption (**Figure 2D**) of *CUL2* and *KLHDC3* supports the essential role of CRL2^KLHDC3^ in UL49.5-mediated TAP degradation.

CRL E3 ligases are activated by conjugation of the ubiquitin-like modifier NEDD8 to cullins (19). Thus, finding UBA3 (NEDD8-activating enzyme E1 catalytic subunit) among the top hits was consistent with the identification of elongins B/C, CUL2, and KLHDC3. We found that the levels of TAP1 and GFP-TAP2 subunits increased following the addition of the neddylation inhibitor, MLN4924 (28), only in the presence of UL49.5 (**Figures 2F, G**). By contrast, nuclear factor erythroid 2-related factor 2 (NRF2) and β-catenin, well-known cullin substrates (29, 30), were stabilized by MLN4924 regardless of the presence of the viral protein (**Figure 2F**). Together, our data demonstrate that the neddylation-regulated CRL2^KLHDC3^ complex is required for UL49.5-mediated TAP degradation.

### UL49.5 acts as an adaptor that recruits CRL2^KLHDC3^ to TAP

The simplest model to explain the requirement for CRL2^KLHDC3^ in mediating UL49.5-dependent TAP degradation is that UL49.5 acts as an adaptor that recruits CRL2^KLHDC3^ to TAP. To test this hypothesis, we used immunoprecipitation to isolate components of the ternary complex in MJS GFP-TAP2 cells depleted of one of the interacting molecules (**Figure 3**). We found that UL49.5 antibodies co-captured all CRL2 E3 ligase components, including the neddylated form of CUL2, ELOB, ELOC, and RBX1, as well as both TAP subunits in UL49.5-expressing, but not control cells (**Figure 3A**), confirming that UL49.5 forms a ternary complex with both TAP and CRL2. Moreover, TAP1 antibodies immunoprecipitated CUL2 and RBX1 together with UL49.5 exclusively in UL49.5-expressing cells (**Figure 3B**), confirming that UL49.5 plays a pivotal role in recruiting the CRL2 E3 to TAP. By contrast, the interaction between CRL2 and UL49.5 was unaffected by TAP disruption in the U937 *TAP1/2* DKO (double knockout) cell line (**Figure 3C**), establishing that UL49.5 binds CRL2 independently of TAP, further supporting the conclusion that this viral protein forms a ternary complex that contains both TAP and CRL2^KLHDC3^.

**Figure 3.**
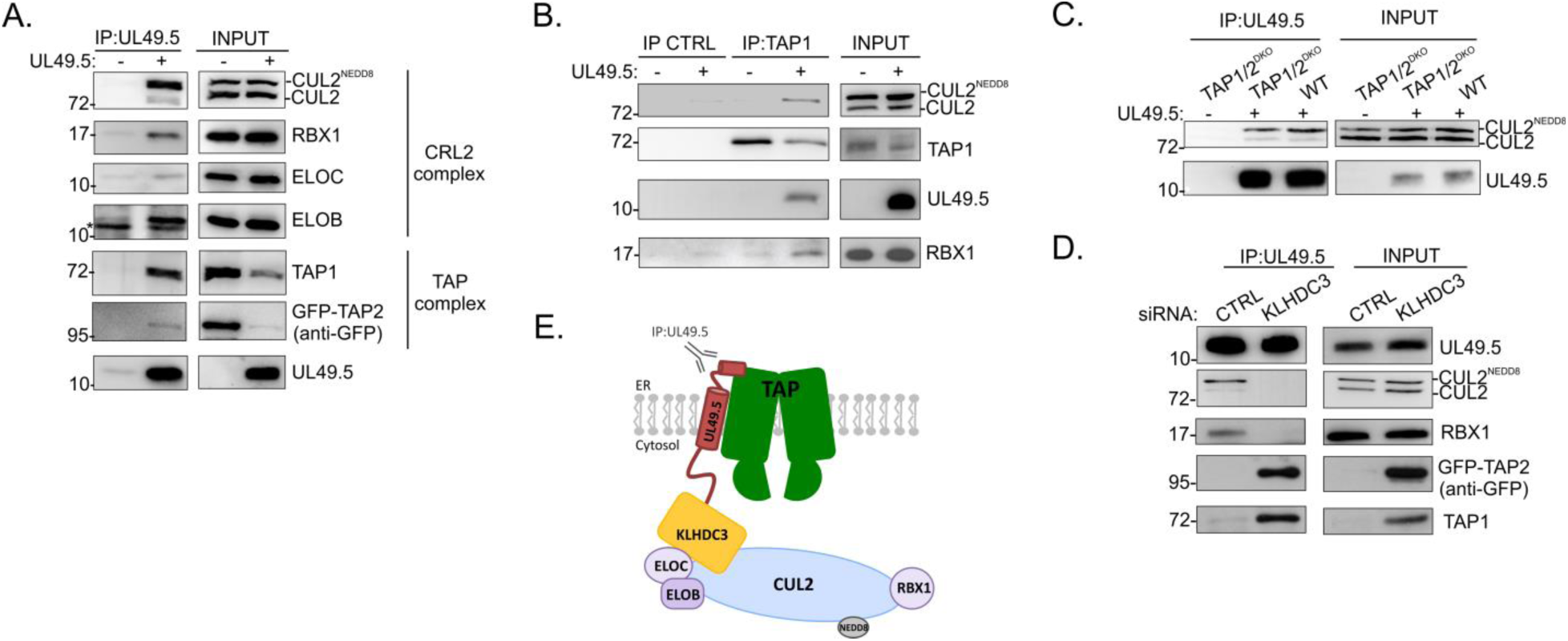
UL49.5 acts as an adaptor recruiting CRL2^KLHDC3^ to TAP. (A) The interaction of UL49.5 with CRL2 and TAP subunits in MJS cells stably expressing UL49.5 was analyzed by UL49.5 IP. * denotes the unspecific ELOB antibody staining. (B) The interaction of TAP1 with CRL2 in MJS cells stably expressing UL49.5 was analyzed by TAP1 IP, followed by immunoblotting. (C) The interaction of UL49.5 with CUL2 in U937 *TAP1/2*^KO^ cells stably expressing UL49.5 was analyzed by UL49.5 IP followed by immunoblotting. (D) MJS cells stably expressing UL49.5 were transfected with siRNA targeting *KLHDC3*. The interaction of UL49.5 with CRL2 and TAP1 was analyzed by UL49.5 IP, followed by immunoblotting. (E) Schematic of UL49.5-TAP-CRL2 ternary complex based on the IP results (the position of the UL49.5-precipitating antibody is marked).

To test the hypothesis that KLHDC3 serves as the SRR linking CRL2 to the UL49.5-TAP complex, we assessed the impact of KLHDC3 depletion on ternary complex formation. While CUL2 and RBX1 were robustly captured with UL49.5 antibodies in siRNA control cells, neither protein was detected in cells treated with *KLHDC3* siRNA. Instead, we observed strongly increased levels of TAP1 and TAP2 subunits, reflecting impaired TAP turnover (**Figure 3D**). These data indicate that UL49.5 interacts with CRL2^KLHDC3^ independently of TAP, yet binding of TAP to this E3 ligase occurs only in the presence of UL49.5 (**Figure 3E**).

### UL49.5 binds CRL2^KLHDC3^ via a C-terminal RGRG degron

To elucidate the mechanism by which UL49.5 recruits CRL2 to TAP, we sought to identify the degron that is recognized by KLHDC3. Previous studies (7,8,31) demonstrated that the cytosolic tail of UL49.5 is essential for TAP degradation, suggesting that the relevant degron resides in this domain. Indeed, antibodies to UL49.5 co-immunoprecipitated CUL2 from cells expressing full-length UL49.5, but not from cells expressing UL49.5Δtail (**Figure 4A**). Moreover, the CUL2-UL49.5 interaction was unaffected by treatment of the cells with MLN4924, indicating that the interaction occurs independently of cullin neddylation.

**Figure 4.**
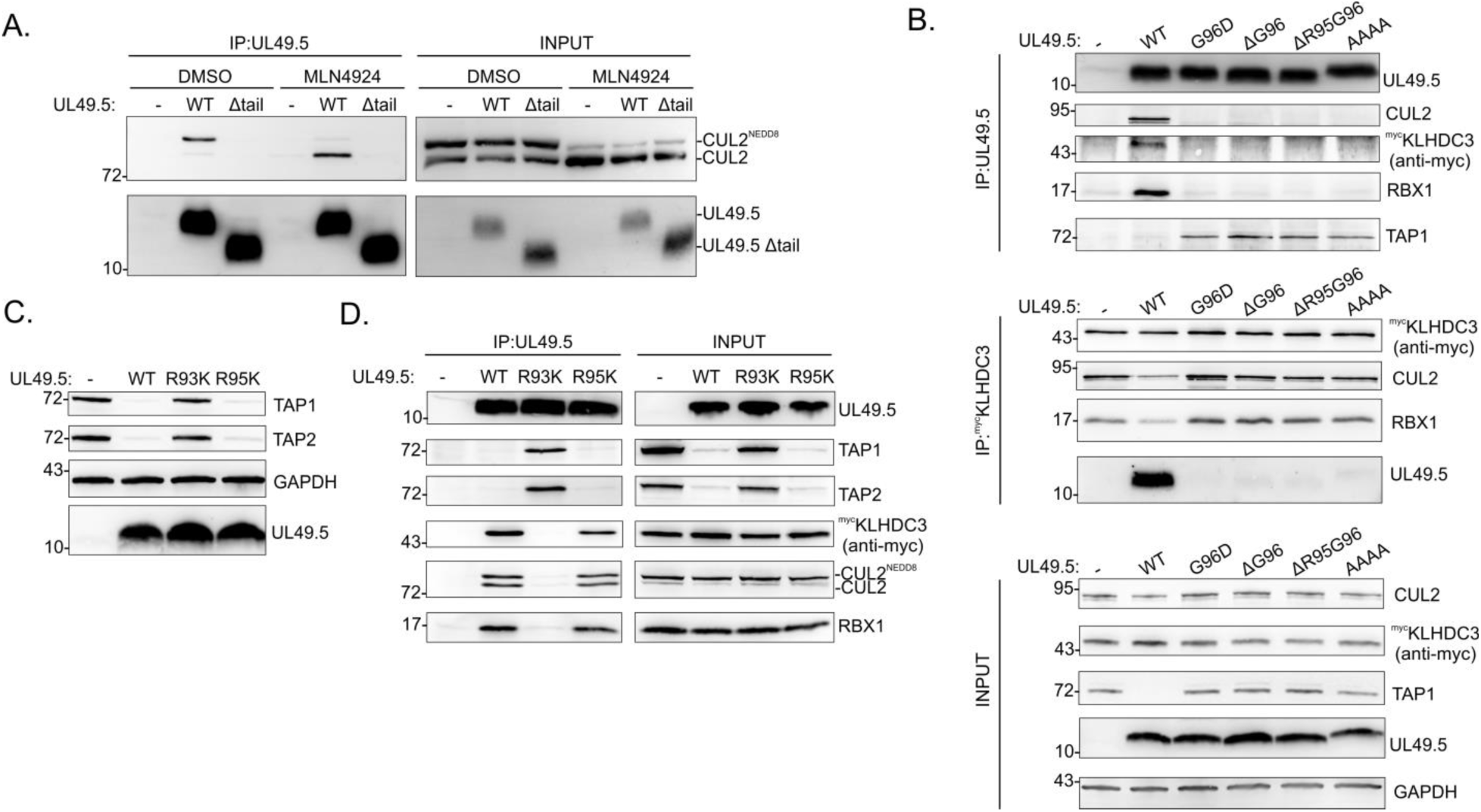
RGRG C-terminal motif of UL49.5 recruits the CRL2^KLHDC3^ complex. (A) MJS cells expressing UL49.5 WT or the UL49.5Δtail were treated for 12 h with MLN4924 or DMSO. The interaction of UL49.5 with CUL2 was analyzed by UL49.5 IP followed by immunoblotting. (B) The interaction of UL49.5 with CRL2 and TAP1 in MJS cells stably expressing ^myc^KLHDC3 and UL49.5 variants was analyzed by ^myc^KLHDC3 and UL49.5 IP followed by immunoblotting. (C) Immunoblotting of steady-state levels of TAP1 and TAP2 in MJS cells stably expressing UL49.5 WT, R93K, or R95K variants. (D) The interaction of UL49.5 WT, R93K, or R95K variants with the CRL2 complex in MJS cells stably expressing ^myc^KLHDC3 was analyzed by UL49.5 IP followed by immunoblotting with the indicated antibodies.

To identify the specific sequence of UL49.5 responsible for the interaction with KLHDC3, we focused on the four C-terminal residues, RGRG^96^, which have been previously shown to be essential for TAP degradation (9). Strikingly, this sequence matches the R(x)nR/KG consensus previously identified as a C-degron that binds directly to KLHDC3 and acts as a degron that recruits CRL2 to its substrate (32–34). To test the hypothesis that the C-terminal RGRG motif in UL49.5 functions as a C-terminal degron that promotes CRL2^KLHDC3^ binding and TAP degradation, we assessed the impact of mutations of the residues comprising this motif. We found that deletion of one (DG96) or two (DR95G96) residues from the C-terminus of UL49.5; substitution of four C-terminal residues with alanine; or the G96D substitution abolished both UL49.5 induced TAP degradation and CRL2^KLHDC3^ binding without affecting the ability of KLHDC3 to bind to CUL2 (**Figure 4B**). The R93K substitution also abrogated both CUL2 association and TAP degradation. By contrast, mutation of the penultimate arginine to lysine (R95K) had no effect on CUL2 association and TAP degradation (**Figure 4C, D**).

In summary, these results point toward the critical role of the arginine residue in position -4, the positively charged residue in position -2, and the C-terminal glycine, to promote the recruitment of KLHDC3, and strongly support the conclusion that UL49.5 promotes TAP degradation by hijacking the host cell C-degron machinery.

### CRL2-recruiting activity of UL49.5 is conserved among human, bovine and murine systems

BoHV-1 UL49.5 is considered a universal TAP inhibitor because it targets and triggers the degradation of not only bovine but also murine and human TAP (35). The mechanism of degron recognition by KLHDC proteins was proposed to be evolutionarily conserved based on the high conservation of KLHDC2 from ameba to humans (36). Therefore, we asked whether CRL2 binding to UL49.5 is a phenomenon of human C-degron machinery, or if it is also conserved. Taking advantage of the fact that bovine cullin 2 could be recognized by anti-human CUL2 antibodies, we performed UL49.5 pulldown from bovine MDBK cells expressing this viral protein and confirmed that UL49.5 forms a complex with CUL2 of bovine origin (**Figure 5A**). Because BoHV-1 UL49.5 has been also reported to cause degradation of mouse TAP in murine colon carcinoma cells (35), we examined UL49.5-CUL2 interaction in the mouse L2 cell line and found that UL49.5 forms a complex also with murine CUL2 (**Figure 5B**). This indicates that hijacking of CRL2 by BoHV-1 UL49.5 for TAP degradation is conserved between species.

**Figure 5.**
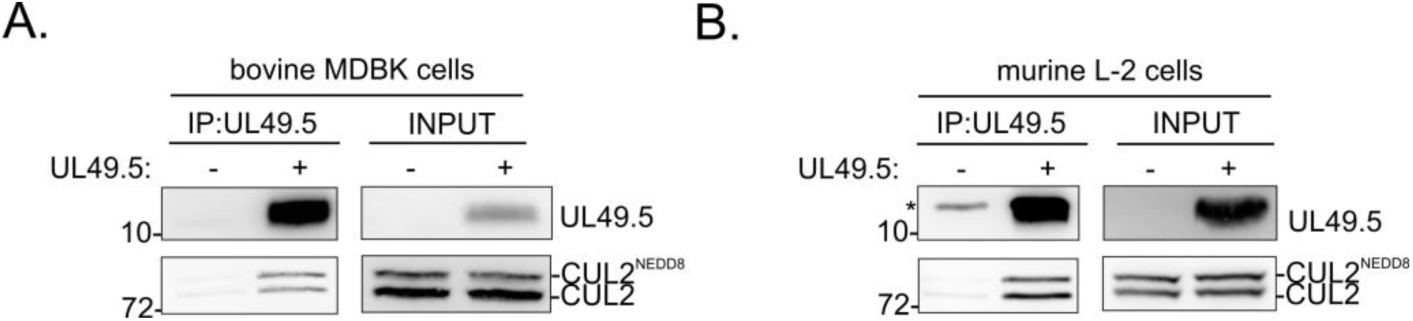
UL49.5 interacts with cullin 2 of bovine and murine origin. (A and B) The interaction of UL49.5 with (A) bovine CUL2 in MDBK cells and (B) murine CUL2 in L2 cells stably expressing UL49.5 was analyzed by UL49.5 IP followed by immunoblotting; *unspecific staining.

### UL49.5 escapes degradation by CRL2^KLHDC3^ despite carrying a C-degron

UL49.5 is itself an unstable protein, exhibiting a half-life in MJS cells of approximately 1 h (**Figure S1)**. Our finding showing that the UL49.5 C-terminal RGRG motif is recognized and bound by KLHDC3 to form a complex with CUL2, suggested that this degron could also promote degradation of UL49.5 independently of its interaction with TAP. Surprisingly, we found that UL49.5 turnover in MJS cells was unaffected by *KLHDC3* knockout (**Figure 6A**). Because UL49.5 is a type I integral membrane protein, we tested whether UL49.5 can be regulated by the classical ERAD HRD1/SEL1L pathway. We found that UL49.5 was strongly stabilized in MJS GFP-TAP2 UL49.5 cells depleted of HRD1 (*SYVN1*), SEL1L, or derlin 2 (*DERL2*) (**Figure 6B** and **6C**). At the same time, *SYVN1*, *SEL1L*, or *DERL2* knockout had no significant impact on the TAP1 level (**Figure 6C**). We observed an increased TAP1 only upon KLHDC3 depletion and accumulation of UL49.5 only in the absence of the HRD1 complex, indicating that UL49.5 and TAP1 are degraded through independent mechanisms (**Figure 6C**).

**Figure 6.**
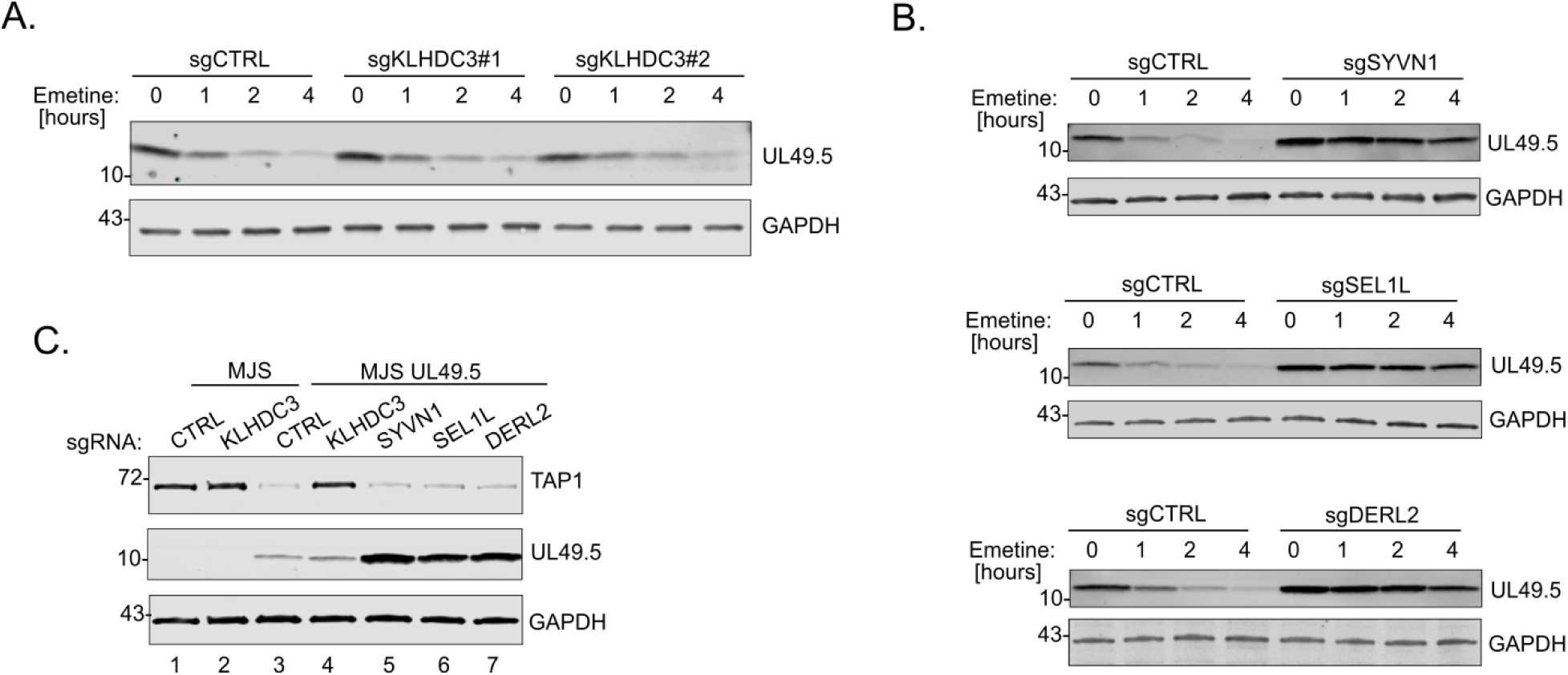
UL49.5 is degraded independently from TAP. (A and B) Turnover of UL49.5 in MJS cells transduced with sgKLHDC3, sgSYVN1, sgSEL1L, or sgDERL2 was analyzed during emetine chase for the indicated times. The level of UL49.5 was detected by immunoblotting. (C) Steady-state levels of TAP1 and UL49.5 in MJS cells transduced with the indicated sgRNAs were analyzed by immunoblotting.

### UL49.5 interacts with CRL2^KLHDC3^ during an early phase of BoHV-1 infection

UL49.5 is known for its dual role during viral infection. In addition to the immunomodulatory, TAP-targeting function, it also forms a complex with BoHV-1-encoded glycoprotein M (gM), which is essential for gM maturation, N-glycosylation, and subsequent virion incorporation (37). These two activities of UL49.5 are regulated in a time-dependent manner during virus infection, dictated by the herpesviral gene expression cascade. Since UL49.5 is an early gene product, it is found in a complex with TAP at the early step of infection. Later, when the late gene product gM is produced, UL49.5 forms a complex with gM that is independent from the TAP ternary complex and traffics to the cell membrane, thereby precluding TAP binding (31). Because of the excess of UL49.5 over gM, there is still a pool of TAP-bound UL49.5 in addition to gM-bound UL49.5 even at the late stage of infection (37).

Knowing that CRL2^KLHDC3^ can interact with UL49.5 in the absence of TAP, we were interested in whether the regulatory effect of gM is an outcome of competition with CUL2 for binding to UL49.5 or if these two proteins can bind UL49.5 independently. Therefore, we analyzed proteins co-immunoprecipitated with UL49.5 at the early (6 h) and late (24 h) time points of BoHV-1 infection in MJS cells. At the early step of infection, when gM was still undetectable by immunoblotting, UL49.5 could be found in a complex with CRL2^KLHDC3^ and TAP subunits; however later, UL49.5 bound predominantly gM, while the amounts of co-immunoprecipitated CUL2, KLHDC3 and RBX1 were reduced (**Figure 7A**). TAP depletion was observed already at the early stage of BoHV-1 infection, and TAP levels were almost undetectable late during infection. We also observed a reduced level of KLHDC3 after 24 hours of infection (**Figure 7A**). We hypothesized that this might result from the combined effect of virus host shutoff endonuclease (vhs, UL41 gene product)-mediated mRNA decay (38, 39) and the short half-life of KLHDC3 of 3 hours (25). To support this hypothesis, we analyzed KLHDC3 expression during infection with the BoHV-1 vhs deletion mutant and found significantly more KLHDC3 in the absence of functional endonuclease (**Figure S3**), suggesting that KLHDC3 expression is repressed over the course of infection.

**Figure 7.**
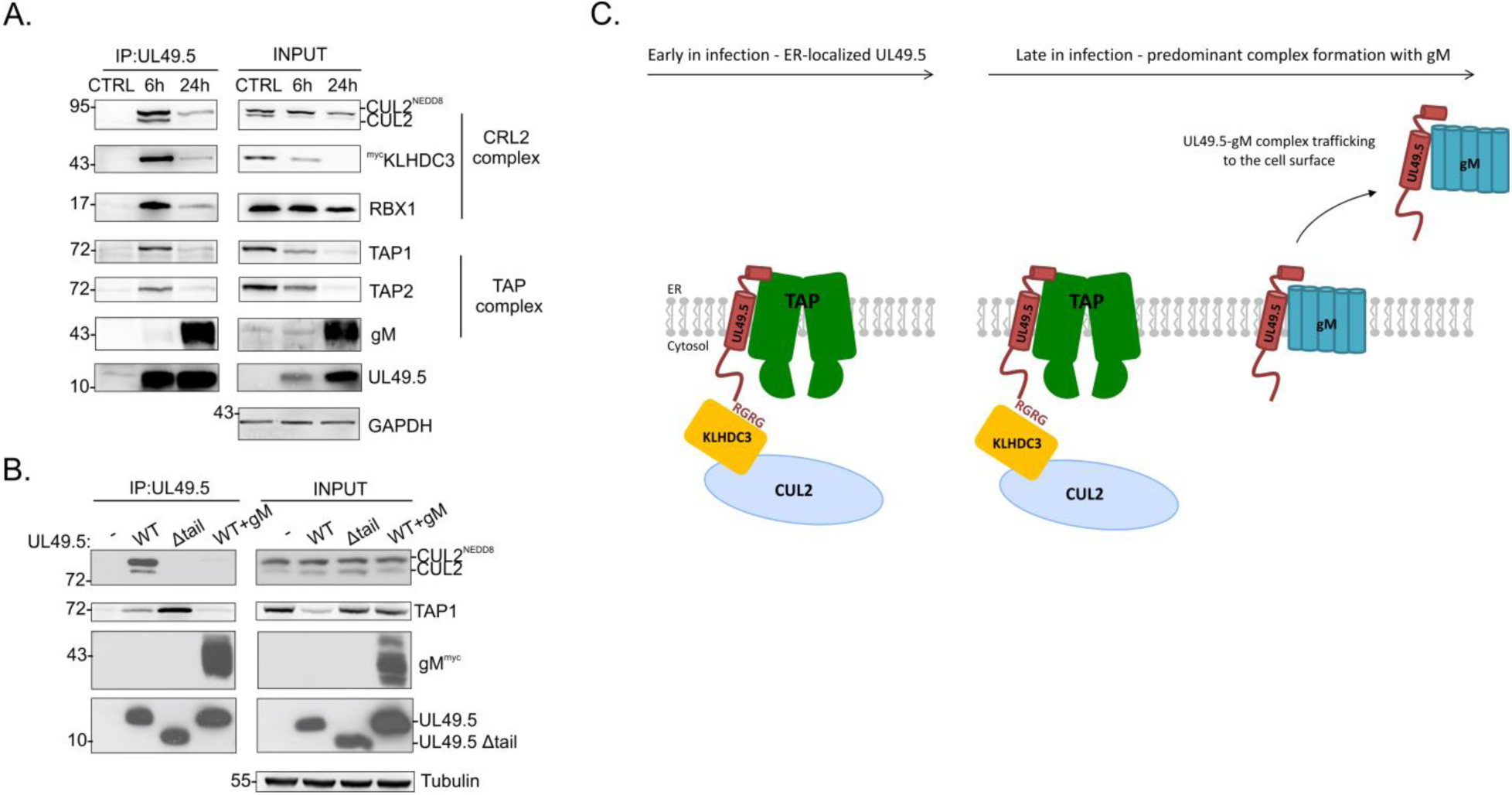
Interaction between UL49.5 and cullin 2 is time-regulated during BoHV-1 infection. (A) MJS ^myc^KLHDC3 cells were infected with BoHV-1 at an MOI of 10 for 6 or 24 hours. The interaction of UL49.5 with CRL2^KLHDC3^, TAP subunits, and gM was analyzed by UL49.5 IP followed by immunoblotting. (B) MJS cells expressing UL49.5 WT, UL49.5Δtail, or UL49.5WT with gM (WT+gM) were subjected to UL49.5 IP followed by immunoblotting. (C) Schematic of time-dependent UL49.5 complex formation during BoHV-1 infection.

To determine if UL49.5 interacts preferentially with gM late in infection because of its higher affinity to gM or because of reduced KLHDC3 levels, we analyzed UL49.5-containing complexes in stable cell lines constitutively expressing viral proteins where the level of KLHDC3 should not be affected. We compared the ability of UL49.5 to interact with CUL2 in cells expressing UL49.5 alone with cells co-expressing UL49.5 and gM. As a CUL2-noninteracting control, we used the Δtail variant of UL49.5. gM co-expression drastically reduced the amount of CUL2 co-immunoprecipitated with UL49.5 (**Figure 7B**), which implies that gM and CRL2 are competing for the interaction with UL49.5. Furthermore, the TAP1 level in cells expressing UL49.5 WT was downregulated, while in cells expressing both UL49.5 WT and gM or UL49.5Δtail, the level of TAP1 was restored. We observed that TAP1 co-immunoprecipitated with UL49.5Δtail, which lacks the CRL2^KLHDC3^-recruiting activity, as well as with UL49.5 WT, however, in reduced amounts due to its degradation. In the presence of gM, only a small portion of TAP1 could be detected in a complex with the UL49.5, which is consistent with the previous report (37).

Taken together, these observations illustrate that TAP is degraded at the early stage of BoHV-1 infection. The interaction between UL49.5 and CRL2 is time-regulated during infection and depends on the presence of gM, which competes with cullin 2 for UL49.5 binding (**Figure 7C**).

## DISCUSSION

Using two complementary genetic screening approaches, we identified the long-sought mechanism employed by the BoHV-1 UL49.5 viral inhibitor to target TAP to ERAD. Unlike herpesvirus proteins mK3 and pK3, which promote the degradation of TAP by functioning as ubiquitin E3 ligases, UL49.5 hijacks a host cell cullin ring ligase via C-degron mimicry to promote TAP degradation by the ERAD.

Identification of CRL2 as an E3 for TAP is unusual since the majority of E3s involved in ERAD are membrane-embedded or membrane-bound enzymes (40), whereas cullins are cytosolic. Only two cases of cullin activity in ERAD have been reported so far: 1) Vpu protein encoded by human immunodeficiency virus 1 exploits cullin 1 (SCF^β-TrCP^) to target CD4 for ERAD (41) and 2) SCF^Fbs1^ and SCF^Fbs2^ were shown to contribute to ERAD by sensing the exposed N-glycans of glycoprotein substrates (42). Our data add to this short list, providing compelling evidence of the involvement of CUL2 in ERAD.

Our results highlight a critical role of BoHV-1 UL49.5’s C-terminal RGRG motif in recruiting CRL2^KLHDC3^ to the TAP complex, explaining why TAP degradation depends on this region of UL49.5 (9). We used systematic mutagenesis to identify a critical role of arginine R93 and the C-terminal glycine (G96) of UL49.5. Previous studies (9) reported that G96 can be replaced by alanine but not with the bulky charged aspartate, as confirmed in this study and in agreement with the the known properties of KLHDC3-dependent C-degrons (32–34). Comparison with the structural details of the diglycine recognition by KLHDC2 (36) suggests that the C-terminal carboxyl group of UL49.5 binds within the KLHDC3 binding pocket. The cryo-EM structure of KLHDC3 would be beneficial to ultimately illustrate KLHDC3-UL49.5 interactions.

Our data add the UL49.5-TAP complex to a short list of KLHDC3 substrates, comprised of UGA-terminated selenoproteins, SEPHS2 and SELS (43), testis-specific Y-encoded-like protein 1 (TSPYL1) (32), and p14ARF (25, 43), identified to date. UL49.5 is unique within this group because, although it shares a consensus C-terminal RGRG motif, it itself escapes KLHDC3-mediated degradation, even though it possesses a consensus degron sequence. Because it is not degraded by KLHDC3, it might be considered a “pseudosubstrate” (44). However, in the context of BoHV-1 infection, UL49.5 acts as an adaptor for a *bona fide* substrate – TAP, thereby functioning as a “trans-degron”, a degron acting *in trans* by proximity-induced degradation. Such degrons, under specific conditions, can recruit E3 ligases to their interacting proteins. A good example of a naturally-occurring *in-trans* degron engagement for the stability regulation are Sc and Da proneuronal development proteins from the bHLH family in *Drosophila melanogaster*, where one protein from a complex provides a degron to ubiquitinate the other, but not for its own removal (45).

It is possible that the potential ubiquitin acceptor sites in TAP become available in structurally distorted, conformationally-arrested TAP and the ones in UL49.5 become inaccessible to CRL2 because of steric hindrance or low exposure from the membrane. Alternatively, ubiquitin-acceptor sites in UL49.5 could be hidden within the KLHDC3 binding pocket. This explains why the half-life of UL49.5 does not depend on KLHDC3, but instead, it is dependent on the HRD1 complex. Degradation of excess UL49.5 by HRD1 might be a self-tuning mechanism, similar to the one described for the US11 protein of HCMV, which triggers MHC I degradation via TMEM129, while excess free US11 is degraded by HRD1 (46).

Interestingly, the comparison of UL49.5 protein sequence from the *Varicelloviruses* revealed that only BoHV-1 and closely related BoHV-5, bubaline herpesvirus 1 (BuHV-1), and cervid herpesvirus 1 (CvHV-1), all of which infect ruminants, encode the C-terminal RGRG motif, which stays in line with their unique ability to induce TAP degradation (9, 16). This observation raises a question about molecular evolution events that resulted in a gain of the RGRG motif.

In the early phase of BoHV-1 infection, recruitment of the CRL2^KLHDC3^ to TAP leads to its degradation. At the late stages of virus infection, the short half-life of KLHDC3 together with vhs-mediated translational host shut-off may limit efficient TAP degradation. Our results demonstrate that UL49.5-encoded C-degron is unique – UL49.5 does not only escape from its own degradation, but it also uses the degron to hijack host ERAD, which contributes to viral immune evasion.

Our data identify many and the most essential components of the cellular machinery that is brought by UL49.5 to TAP for its degradation. However, the role of factors, like FAF2 or BAG6 requires further experimental validation. The remaining questions include temporal-spatial details of UL49.5-KLHDC3-TAP complex formation. We demonstrate that UL49.5-KLHDC3 binding is TAP-independent whereas TAP forms a complex with CRL2 only in the presence of the viral protein, but it is not fully clear what is the sequence of these events, e.g., does UL49.5 bind KLHDC3 or TAP first or if UL49.5 is recognized by HRD1 before or after KLHDC3 dissociation.

The biological significance of our findings revolves not only around explaining the molecular mechanism of TAP degradation but has some potential therapeutic applications. Conservation of TAP degradation for human, bovine, and murine TAP, has already brought attention to its use in herpes simplex-based oncolytic T-stealth vectors for anticancer therapies, where HSV-1-encoded TAP inhibiting protein ICP47 was replaced with more efficient BoHV-1-encoded UL49.5 (47). UL49.5 may also serve as a biological tool to study the C-degron pathway and the mechanism of substrate recognition by KLHDC3, which may contribute to targeted protein degradation and the rational design of PROTACs (48).

## ACKNOWLEDGMENTS

We thank Rajat Rohatgi and Mandi Ma from the Departments of Biochemistry and Medicine, Stanford University School of Medicine for sharing the Brunello library. We are grateful to Julien Couthouis, for help with analysis of the CRISPR/Cas9 screen data. We would like to thank Alicja Sznarkowska from the International Centre for Cancer Vaccines Science, University of Gdańsk, for sharing NRF2 reagents and Katarzyna Rückemann-Dziurdzińska from the Division of Pathology and Experimental Rheumatology, Medical University of Gdańsk, for the assistance with cell sorting, Piotr Brągoszewski from Nencki Institute for Experimental Biology, for help in designing the UPS siRNA library. We thank Samantha C. Gumbin and Francesco Scavone from the Department of Biology, Stanford University for critical reading of the manuscript.

## AUTHOR CONTRIBUTIONS

Conceptualization, M.W., C.R., R.J.S, R.R.K., and A.D.L; Methodology, M.W., C.R.,, M.G., L.S.B., W.Q., R.J.S., R.R.K., and A.D.L.; Formal Analysis, M.W., C.R., L.S.B., W.Q., R.J.S., and A.D.L.; Investigation, M.W., C.R., M.G., M.M. and A.D.L.; Resources, J.E.C., K.B-S., R.J.S., R.R.K., and A.D.L.; Writing – Original Draft, M.W., R.R.K., M.J.Ś., and A.D.L; Writing – Review & Editing, M.W., C.R., R.J.S., R.R.K. and A.D.L., Visualization, M.W., C.R., and L.S.B; Supervision, R.J.S., R.R.K., and A.D.L.; Funding Acquisition, M.W., C.R., R.R.K., and A.D.L.; Project administration, M.W. and A.D.L.

## Key resources table

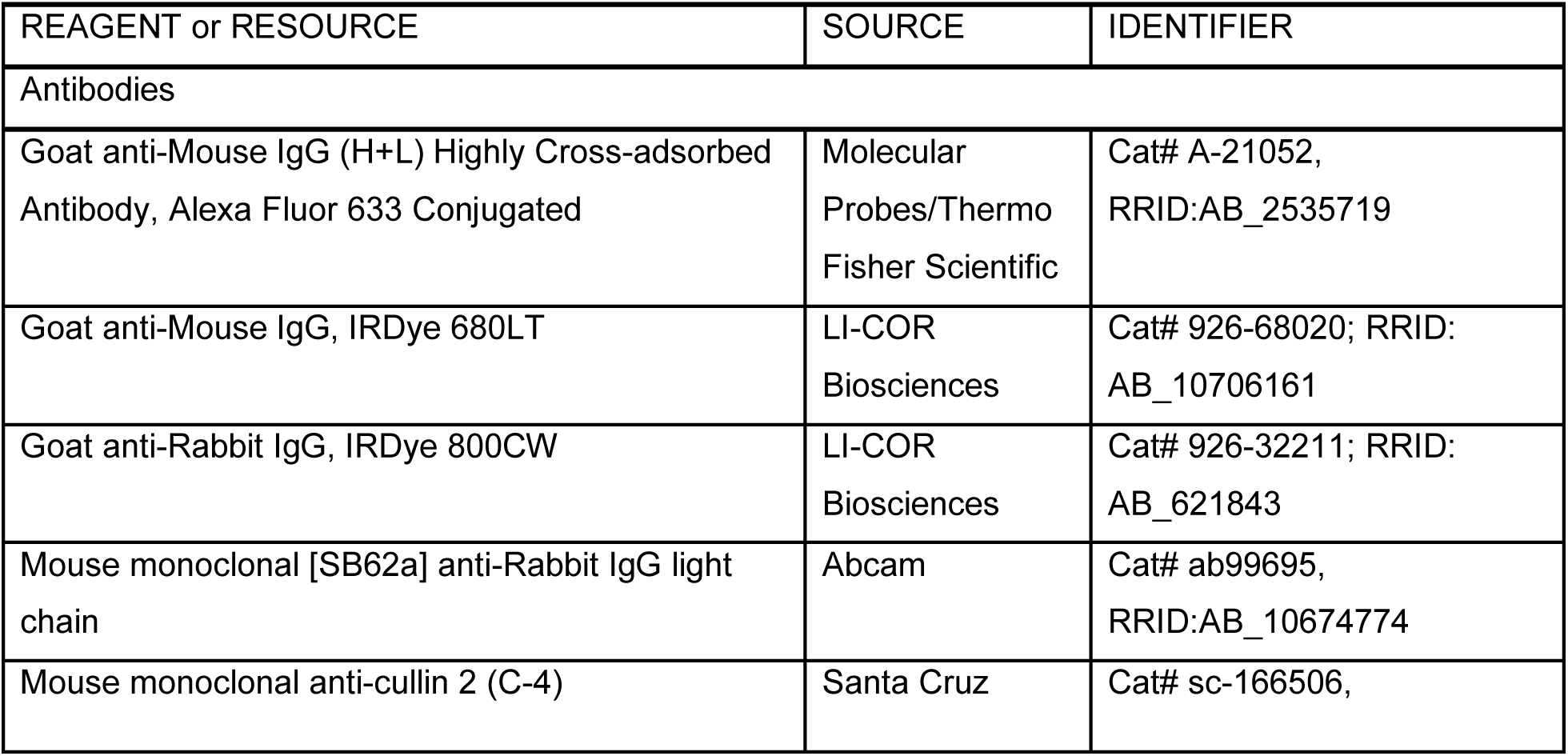
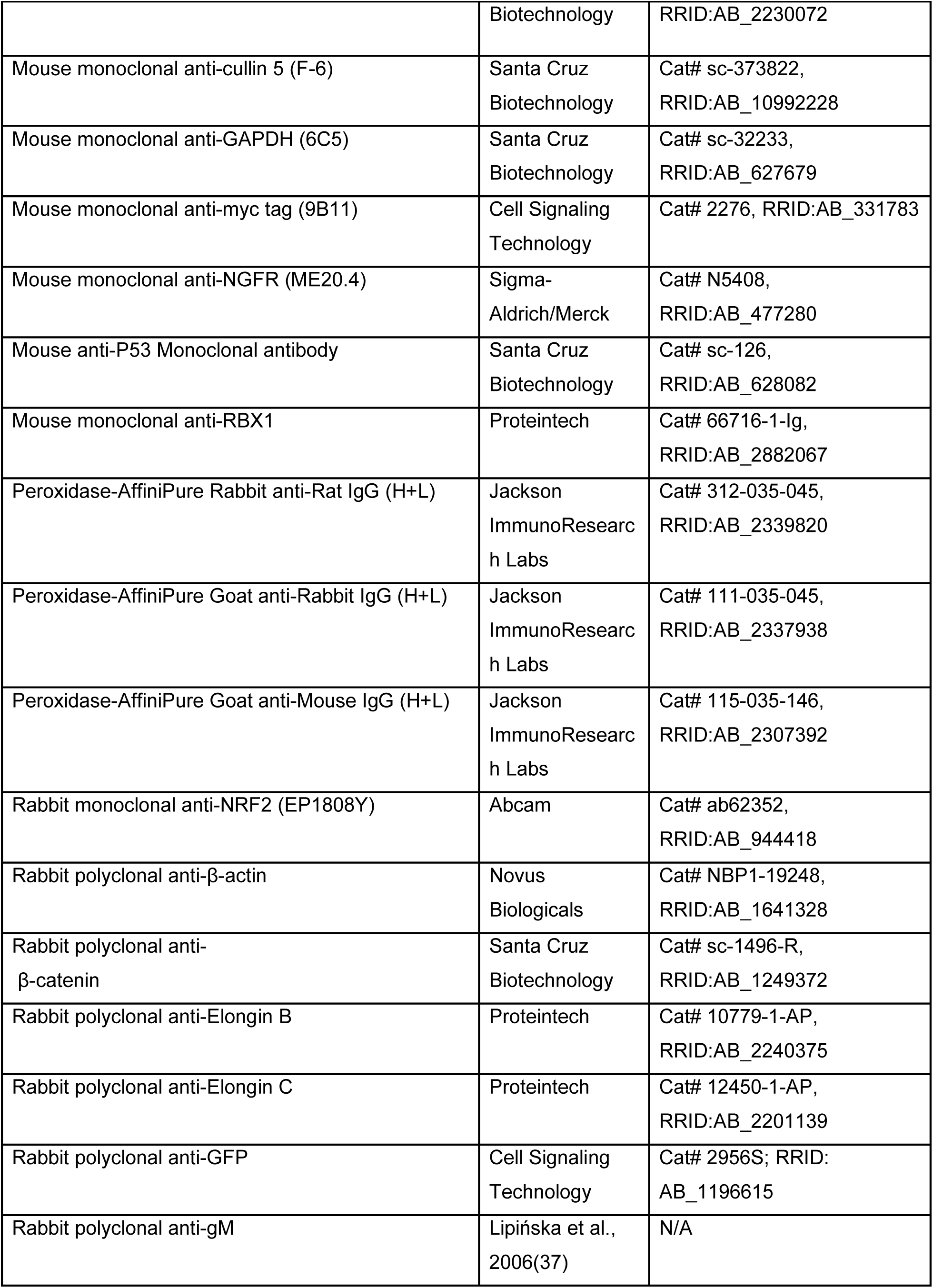
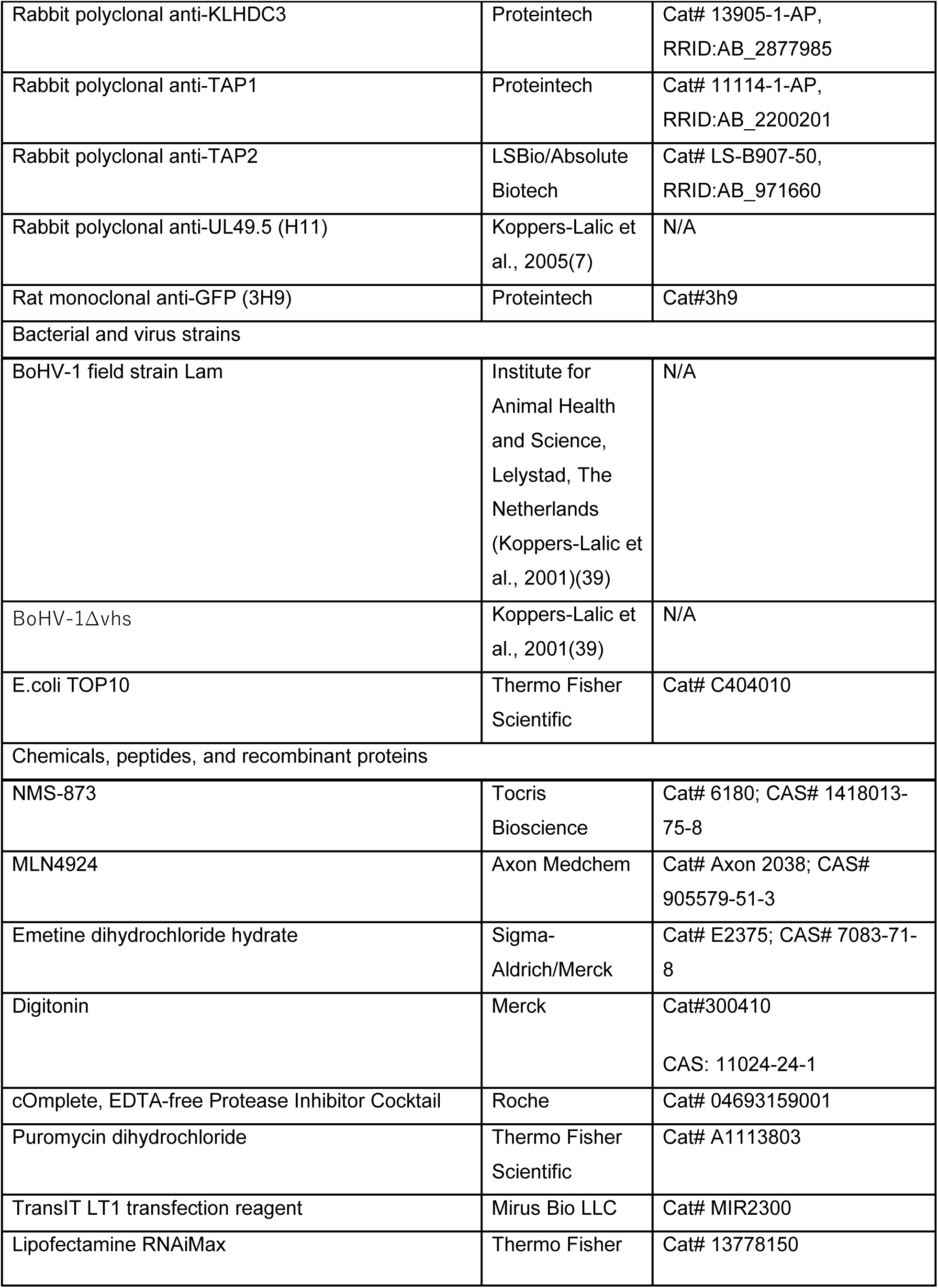
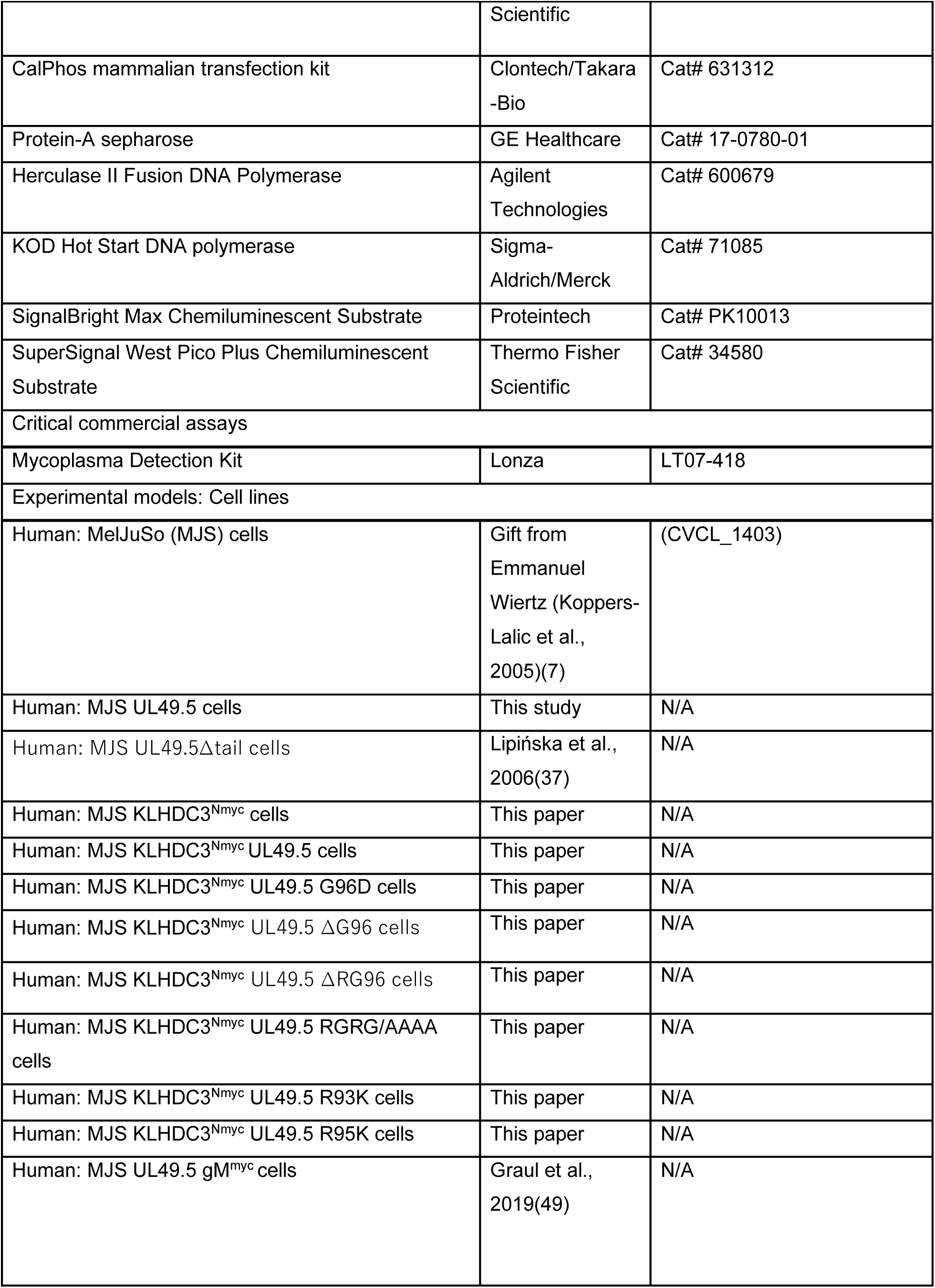
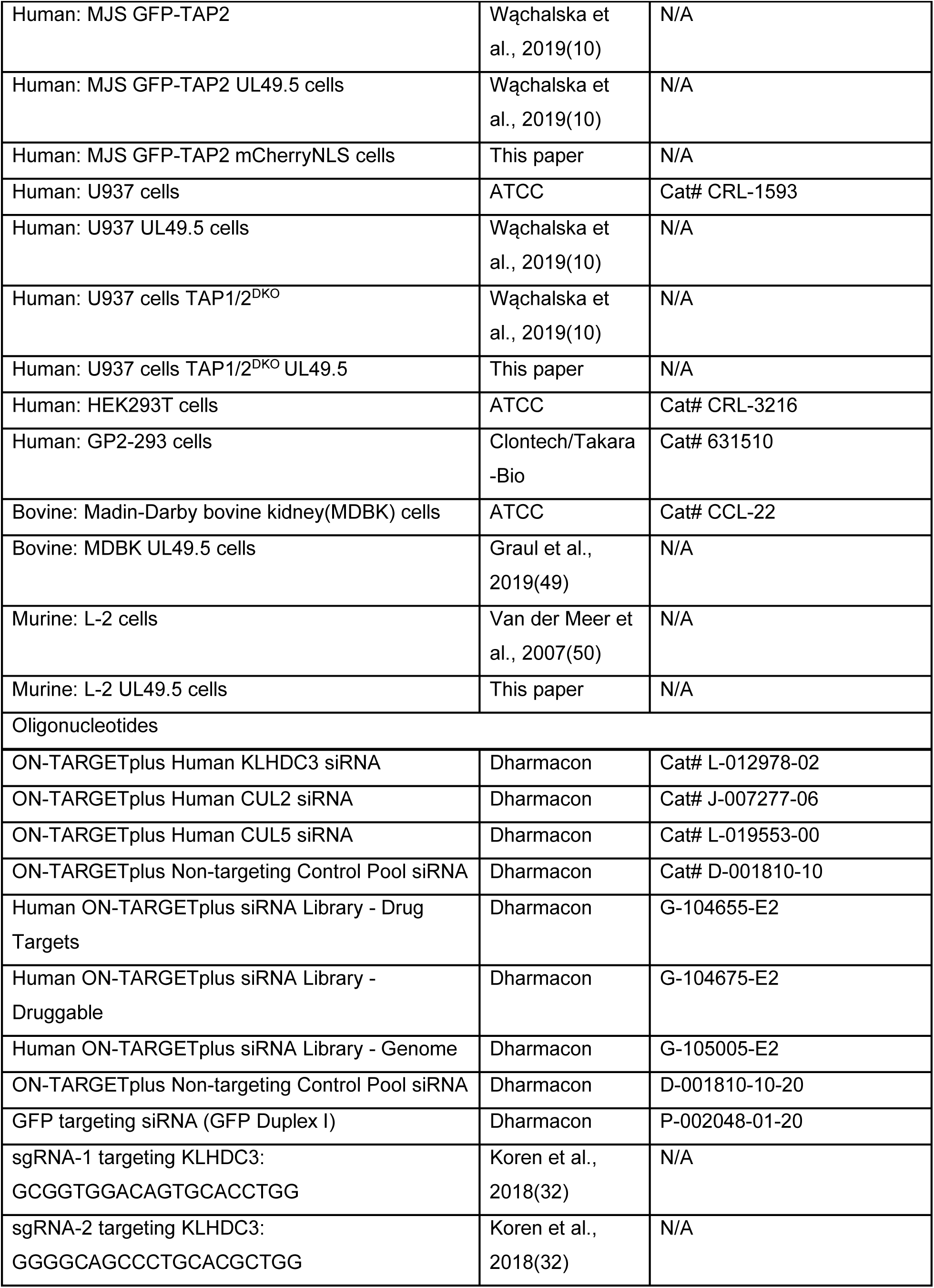
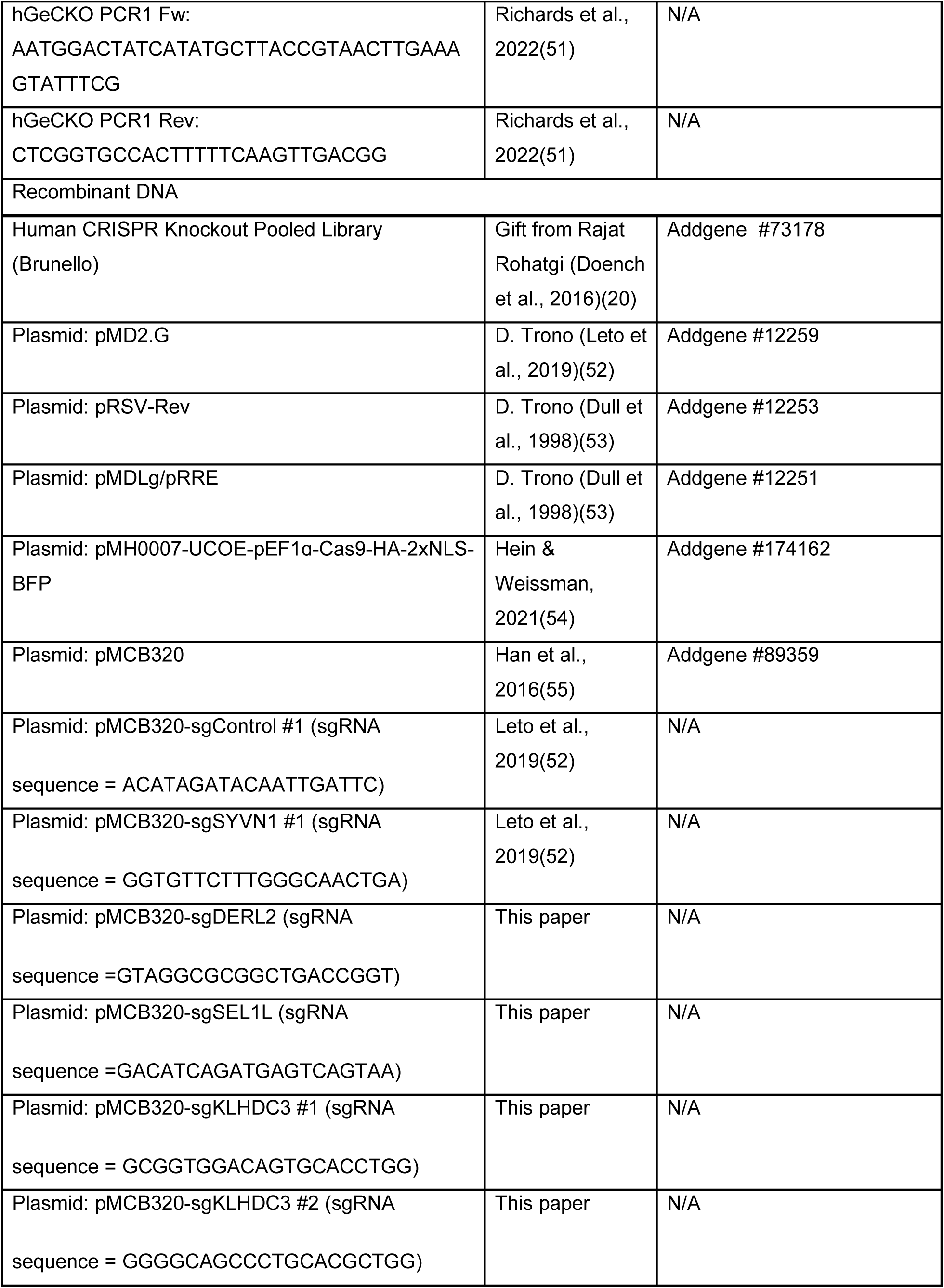
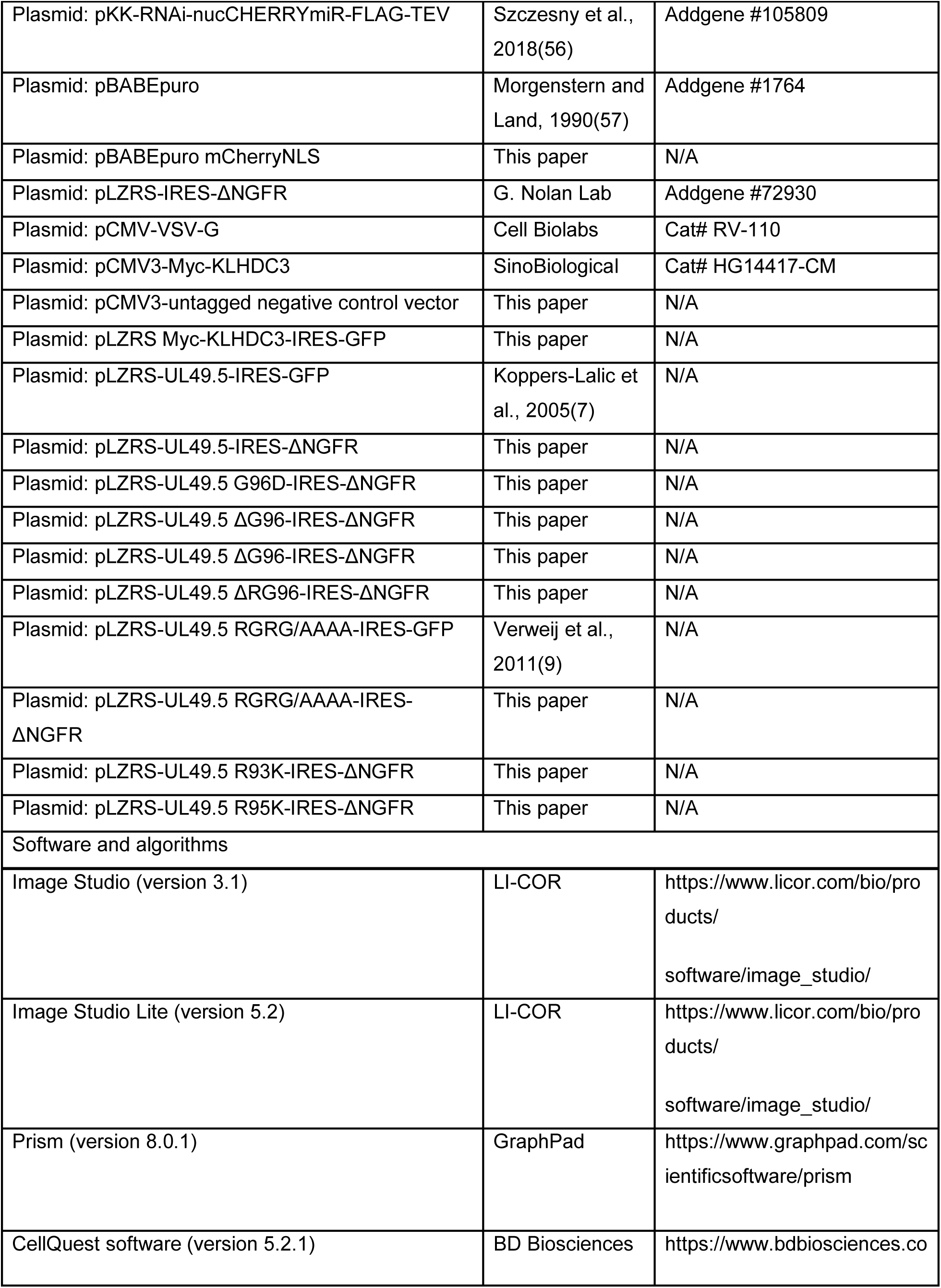
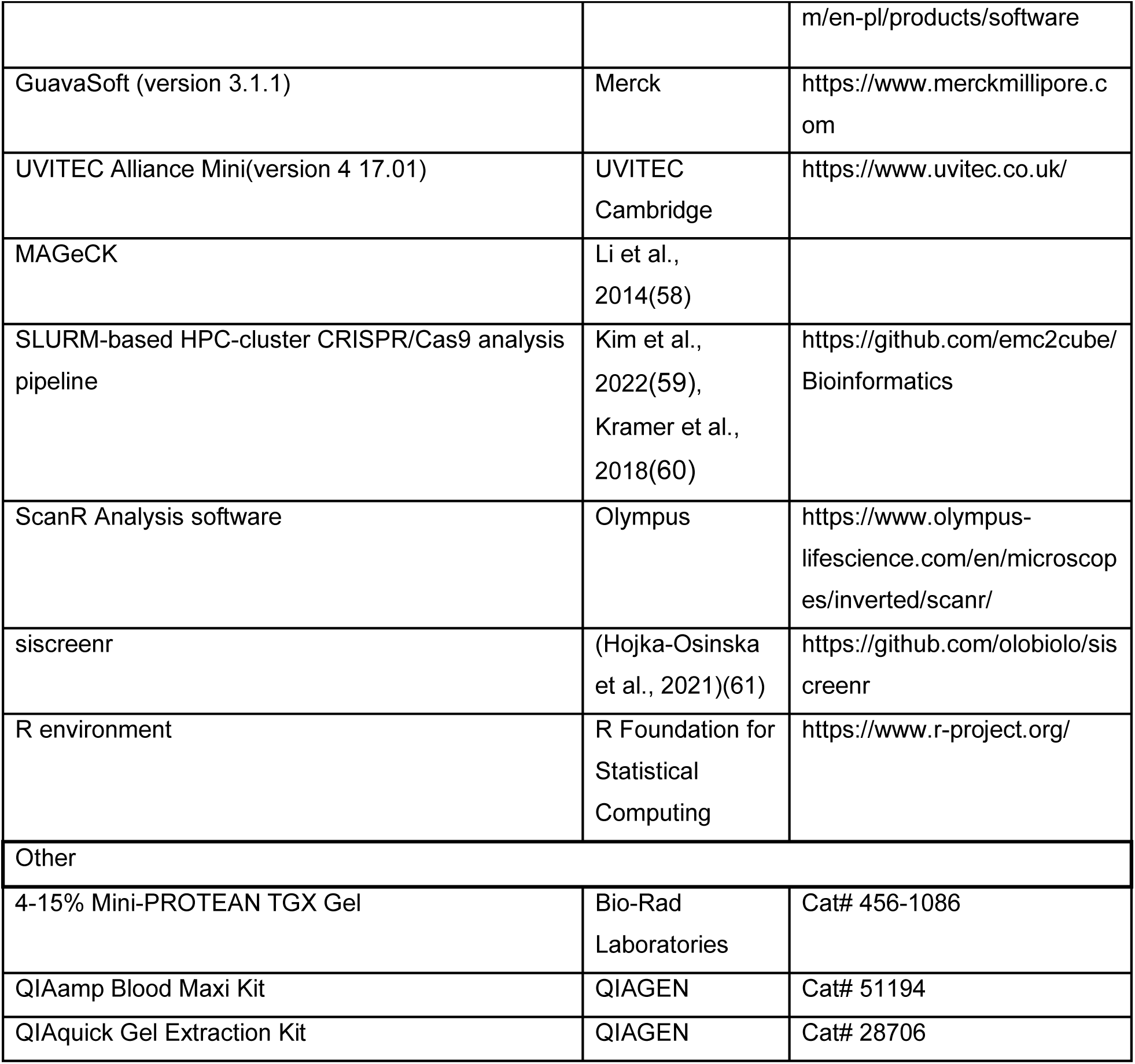

## EXPERIMENTAL MODEL AND SUBJECT DETAILS

### Cell lines

Human melanoma Mel JuSo (MJS) cells, U937 cells (ATCC, CRL-1593), and Madin-Darby bovine kidney (MDBK) cells (ATCC, CCL-22) were cultured at 37°C and 5% CO2 in RPMI 1640 (Corning) supplemented with 10% v/v FBS, 2 mM L-glutamine, 100 IU/ml penicillin, 100 mg/ml streptomycin and 0.25 mg/ml amphotericin B (all from Thermo Fisher Scientific). Murine L-2 cells(50) HEK293T (ATCC, CRL-3216) used for lentivirus production were grown in DMEM (Corning) and GP2-293 cells (Takara/Clontech) used for retrovirus production were cultured in Iscove’s modified Dulbecco’s medium (IMDM, Lonza); supplemented as above. BoHV-1 and its vhs deletion mutant (BoHV-1 vhs-)(39) were propagated and titrated on MDBK cells as described previously(18).

## METHOD DETAILS

### Plasmid construction

pLZRS-UL49.5-IRES-NGFR and pLZRS-UL49.5RGRG/AAAA-IRES-ΔNGFR constructs were generated by subcloning UL49.5 and UL49.5 RGRG/AAAA genes from the pLZRS-UL49.5-IRES-GFP(7) and pLZRS-UL49.5RGRG/AAAA-IRES-GFP(9) vectors, respectively, to the retroviral expression vector pLZRS-IRES-ΔNGFR, upstream of an internal ribosome entry site (IRES) element, which is followed by ΔNGFR (truncated nerve groWTh receptor, used here as a marker gene). Other UL49.5 variants were generated by PCR using KOD Hot Start DNA polymerase (Merck), pLZRS-UL49.5-IRES-GFP(7) as a template, and primers listed in Table S3. PCR products were inserted into pLZRS-IRES-ΔNGFR. The identity of the constructs was verified by DNA sequencing. pBABEpuro mCherryNLS was obtained by subcloning of nucmCherry from pKK-RNAi-nucCHERRYmiR-FLAG-TEV(56) to the pBABEpuro vector. pLZRS Myc-KLHDC3-IRES-GFP construct was generated by subcloning Myc-KLHDC3 gene from pCMV3-Myc-KLHDC3 to the pLZRS-IRES-GFP vector. The pCMV3-untagged negative control vector was obtained by deleting the Myc-KLHDC3 gene and re-ligating the empty vector. To generate sgRNA expression plasmids, pairs of oligonucleotides containing the sgRNA sequence and BstXI and BlpI overhangs were phosphorylated and annealed. Annealed oligos were then ligated into the BstXI and BlpI-digested and gel purified pMCB320 vector backbone(55).

### Retroviral and lentiviral transduction

For recombinant retroviruses, the transfer plasmid (pBABEpuro-based or pLZRS-IRES-ΔNGFR/GFP-based) and pCMV-VSV-G were co-transfected into GP2-293 packaging cells using CalPhos mammalian transfection kit (Takara/Clontech) according to manufacturer’s protocol. 24 hours after the transduction the medium was replaced. Virus-containing supernatants were collected after 48 h, passed through a sterile 0.45 µM syringe PVDF filter (Millipore), and used for transduction in the presence of 8 µg/mL polybrene (Sigma-Aldrich). 24 hours after transduction the medium was replaced and cells were grown in fresh RPMI medium for 48 hr. Transduced cells were selected by growing in RPMI medium containing 2 µg/mL puromycin (Thermo Fisher Scientific) for 3-5 days or sorted for NGFR-positive or GFP-positive cells.

For the production of sgRNA lentiviruses, the pMCB320 plasmid(55) containing the indicated sgRNA was combined with a third-generation lentiviral packaging mix (1:1:1 mix of pMD2.G, pRSV-Rev, and pMDLg/pRRE) at a 1:1 ratio and transfected into HEK293T cells using TransIT LT1 transfection reagent (Mirus Bio LLC). After 72 h virus-containing supernatants were collected, filtered through a 0.45 uM PVDF filter (Genesee Scientific), and cells were transduced as described for retroviruses, followed by puromycin (2 µg/mL) selection for 3-5 days.

MJS KLHDC3^Nmyc^ cells were obtained by transduction with a pLZRS-IRES-GFP-based retrovirus and sorting for GFP-positive cells. UL49.5 variant-expressing cell lines were generated with pLZRS-IRES-NGFR-based retroviruses and sorted for NGFR-positive cells. MJS GFP-TAP2 UL49.5 Cas9-BFP cells were obtained by transduction of MJS GFP-TAP2 UL49.5 cells(10) with the pMH0007-UCOE-pEF1ɑ-Cas9-HA-2xNLS-BFP vector(54) and were sorted two times for BFP-positive cells using a BD FACSAria II Cell Sorter (BD Biosciences) at the Stanford Shared FACS Facility. MJS GFP-TAP2 mCherryNLS cells were generated by a pBABEpuro-based retrovirus transduction of MJS GFP-TAP2 cells(10), followed by puromycin (2 µg/mL) selection.

### Viral infection of cells

MJS cells were washed once with PBS and infected with BoHV-1 at an MOI of 10 in serum-free medium. After 1 h, complete RPMI medium was added. Cells were collected for cell lysis and immunoprecipitation after 6 or 24 hours. In all experiments, mock-infected cells were treated under the same conditions as infected cells.

### Flow cytometry

MJS cells were trypsinized, washed once with PBS, re-suspended in 0.5 ml PBS, and kept on ice until data acquisition. GFP fluorescence was analyzed by the Guava easyCyte flow cytometer (Merck) and GuavaSoft software (version 3.1.1).

Cells were sorted based on GFP or ΔNGFR using a FACS Calibur with sorting option (BD Biosciences) and CellQuest software (5.2.1). For NGFR-based cell sorting, anti-NGFR antibodies (Sigma-Aldrich) and secondary Alexa 633-conjugated antibodies (Molecular Probes) were used. Cas9-BFP expressing cells were sorted on an Aria II cell sorter (BD Biosciences) using the BD FACS Diva software.

### RNAi-mediated knockdown

MJS cells were reverse transfected using RNAiMax (Thermo Fisher Scientific) according to the manufacturer’s instructions. The following ON-TARGETplus siRNAs (Dharmacon) were used: siRNA cullin-2 (J-007277-06), siRNA cullin-5 (L-019553-00), siRNA KLHDC3 (L-012978-02), siRNA non-targeting control pool (D-001810-10). Cells were analyzed 72-96 hours post-transfections.

### Cell lysis and immunoprecipitation

Cells were trypsinized, washed with PBS, and lysed in buffer containing 0.5% NP-40 in 50 mM Tris-HCl pH 7.5, 5 mM MgCl2, or in 1% (w/v) digitonin in 50 mM Tris-HCl pH 7.5, 5 mM MgCl2, 150 mM NaCl, and the Complete mini protease inhibitor cocktail (Roche) for 45 min on ice. Lysates were pre-cleared by centrifugation at 12,000 x g for 20 min at 4°C. For immunoblotting, samples were incubated in Laemmli buffer containing 1,25% (v/v) β-mercaptoethanol for 10 min at 95°C or 20 min at 65°C. For immunoprecipitation, pre-cleared cell lysates were rotated overnight at 4°C with anti-UL49.5 antibodies (H11) or anti-TAP1 antibodies (Proteintech) together with Protein-A Sepharose beads (Merck). Beads were washed 3x in 0.1% (w/v) digitonin in 50 mM Tris-HCl pH 7.5, 5 mM MgCl2, 150 mM NaCl buffer, and proteins were eluted by incubating in Laemmli buffer for 10 min at 95°C or 20 min at 65°C.

### SDS-PAGE and Immunoblotting

Protein lysates were separated by SDS-PAGE and transferred to PVDF or nitrocellulose membranes. Proteins were detected using specific primary antibodies followed by incubation with HRP-conjugated or fluorescent IRDye secondary antibodies. Images were acquired with UVITEC Cambridge Mini HD instrument (UVITEC) using chemiluminescence substrates (SuperSignal West Pico Plus substrate (Thermo Fisher Scientific) or Signal Bright Max Chemiluminescent Substrate (Proteintech) or by Odyssey CLx imaging system (Li-COR Biosciences).

### Emetine chase

MJS cells were treated with 20 µM emetine (Sigma-Aldrich) for the indicated times and harvested for cell lysis as described above. Lysates from equal amounts of cells were analyzed by immunoblotting. Western blot band intensities were quantified using ImageStudio Lite version 5.0.21 (LI-COR Biosciences) and normalized to the loading control. The remaining protein was calculated as a percentage of the initial amount detected at time 0 and one-phase exponential decay curves were fit using Prism 8.0.1 (GraphPad Software).

### Genome-wide CRISPR/Cas9 screen

190 million MJS GFP-TAP2 UL49.5 cells that constitutively express Cas9 were transduced with the CRISPR KO lentiviral Brunello library(20) (a kind gift from Rajat Rohatgi and Mandi Ma) at an MOI of 0.4 to achieve > 1000X coverage of the Brunello library. Cells were selected with puromycin (2 µg/mL) for 7 days and pooled together. Cas9-BFP+ cells were sorted for top 5% of the brightest GFP signal simultaneously on two BD FACSAria II Cell Sorters (BD Biosciences) equipped with blue 488 nm and violet 405 nm lasers. Approximately 3 million cells were collected for each duplicate to ensure 1000X coverage of the Brunello library. Genomic DNA was extracted from both sorted and unsorted control cells using the QIAGEN BloodMaxi kit. sgRNAs were PCR amplified from the genomic DNA using Herculase II Fusion DNA Polymerase (Agilent Technologies) and the flanking primers (hGeCKO_F1 and hGeCKO_R1). Each PCR reaction contained: 6 µg of genomic DNA, 20 µL 5X Herculase buffer, 1 µL of 100 mM dNTPs, 1.5 µL 25 µM hGeCKO_F1, 1.5 µL 25 µM hGeCKO_R1, 2 µL Herculase II Fusion DNA polymerase, and water to adjust the volume to 100 µL. The sgRNAs were amplified using the following PCR protocol: 1x 95°C for 2 min, 16x 95°C for 20 s, 60°C for 20 s, 72°C for 30 s, 1x 72°C for 3 min. PCR1 product was used for the second PCR reaction where Illumina adapters and a barcode to DNA for each individual sample were added (Table S3). PCR reactions were set up for each sample as follows: 5 µL of PCR reaction 1, 20 µL 5X Herculase buffer, 1 µL of 100 mM dNTPs, 2.5 µL 10 µM hGeCKO barcoded Fw and Rev primer, 2 µL Herculase II Fusion DNA polymerase, and water to adjust the volume to 100 µL. The sgRNAs were amplified using the following PCR protocol: 1x 95°C for 2 min, 16x 95°C for 20 s, 60°C for 20 s, 72°C for 30 s, 1x 72°C for 3 min. The PCR products were separated in 2% agarose gels, excised, and purified using a QIAGEN Gel Purification Kit. The sgRNAs were analyzed at Novogene (Sacramento, CA) on an Illumina HiSeq using a paired-end sequencing strategy to obtain 150 base-pair reads. Enrichment of each guide RNA was calculated by comparing the relative abundance in the selected and unselected population using MAGeCK(58) to calculate RRA scores. MAGeCK analysis was performed using a SLURM-based HPC-cluster CRISPR/Cas9 screening analysis pipeline(62, 63) on Sherlock, a high-throughput computing cluster managed by the Stanford Research Computing Center

### Cell transfection for siRNA screen

siRNA handling and transfection were performed similarly as described in Hojka-Osinska et al., 2021(61). The screening was performed in triplicate in 384-well plates (Greiner) with a poly-L-lysine-coated µClear bottom. The siRNA library was stored at − 80°C in 384-well plates. A day before transfection, an aliquot of siRNA stock was transferred to a master plate that was subsequently replicated to transfection plates. The transfection plates contained 5 µl of 140 nM siRNA per well. The plates were then sealed and stored at 4°C overnight. The next day, Lipofectamine RNAiMAX (Thermo Fisher Scientific) was added to the wells by dispensing 10 µl of OptiMEM mixed with 0.1 µl of RNAiMAX. This was performed with a Multidrop Combi dispenser (Thermo Fisher Scientific). MJS GFP-TAP2 mCherryNLS and MJS GFP-TAP2 UL49.5 cells were grown separately. On the day of transfection, they were detached, counted, and mixed at a ratio of 1:1, and 20 µl of cell suspension (850 cells) per well was added to the transfection plates with a Multidrop Combi dispenser. After one hour of incubation at room temperature, the plates were moved to a cell culture incubator. 96 hours after transfection, the cells were fixed and stained with 35 µl of 8% formaldehyde in PBS with 4 µg/ml Hoechst 33342 for 40 min at room temperature. Then, they were washed four times with PBS using a 405 LS Microplate Washer (BioTek) and covered with 50 μl of PBS with 1% pen-strep solution. The plates were sealed and stored at 4°C until imaging, for no more than one week.

### siRNA library generation

The protein homeostasis siRNA library was created by picking 1998 siRNA pools from the Human Genome ON-TARGETplus siRNA Library-SMARTpool (Dharmacon) that covers 18,104 genes and is comprised of three subsets: Drug Targets (G-104655-E2, Lot 11169), Druggable (G-104675-E2, Lot 11167), and Genome (G-105005-E2, Lot 11170). Each transfected plate contained a set of control siRNAs: non-targeting siRNA (ON-TARGETplus Non-targeting Pool, D-001810-10-20, Dharmacon) and GFP-targeting siRNA (GFP Duplex I, P-002048-01-20, Dharmacon).

### Image acquisition and analysis

Imaging of siRNA plates was carried out using a ScanR system (Olympus), which was equipped with an MT20 illumination unit and a 150 W mercury-xenon burner, and a DAPI/FITC/Cy3/Cy5 quad Sedat filter set (Semrock) using a UPLANSAPO 20x/0.75 NA lens (Olympus). Single images (not stacks) were taken for three fluorescence channels (Hoechst, GFP, mCherry) from six fields of view in each well. Analysis of the primary images was performed using ScanR Analysis software. Each channel was subjected to background subtraction using the rolling-ball method. Segmentation was started from the Hoechst (nuclei) channel, using the Edge algorithm to identify the main objects. The identified objects were gated based on a scatter plot of the circularity coefficient against the area. The objects that comprised the gating are hereafter referred to as cells. A fixed distance around the main objects (nuclei) was masked as sub-objects to measure the GFP signal in the cytoplasm. The average intensity of the GFP channel in the sub-objects was calculated for all cells. MJS GFP-TAP2 mCherryNLS and MJS GFP-TAP2 UL49.5 cells were distinguished based on the mCherry signal in the nuclei.

### siRNA screen hit selection

Viability was calculated by comparing the number of cells in a given well with the number of cells transfected with non-targeting siRNA in a given plate. For further analysis, we omitted wells (genes) with less than 10% viability. The ratio of the EGFP channel in MJS GFP-TAP2 UL49.5 to MJS GFP-TAP2 mCherryNLS was calculated, averaged between replicates, log-transformed, and z-score calculated using the R environment.

## QUANTIFICATION AND STATISTICAL ANALYSIS

Flow cytometry data were quantified by calculating the mean fluorescence intensity of the population using GuavaSoft 3.1.1 (Merck). Statistical significance was assessed by one-way ANOVA using Prism version 8.0.1 (GraphPad Software, USA).

---

**Supplementary Figure 1.**
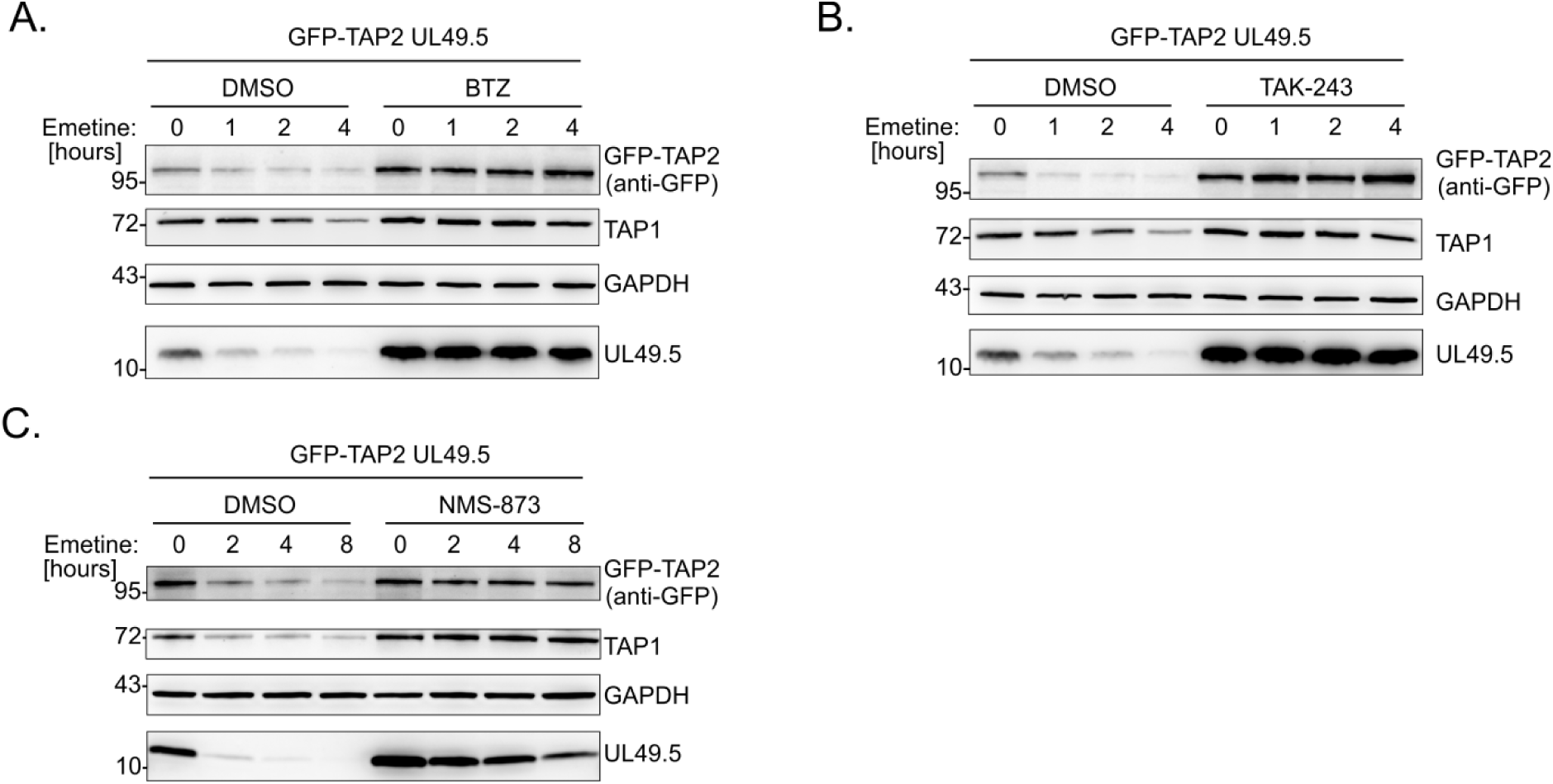
UL49.5-induced TAP degradation depends on ubiquitin-proteasome system and VCP/p7. MJS GFP-TAP2 stably expressing UL49.5 cells were treated with (A) the proteasome inhibitor bortezomib (2 µM) (BTZ) for 4 hours or (B) the ubiquitin-activating enzyme, UBA1, inhibitor TAK-243 (1 µM) for 4 hours, or (C) the VCP/p97 inhibitor NMS-873 (2 µM) for 24 hours. Turnover of TAP1, GFP-TAP2, and UL49.5 was assessed by immunoblotting lysates from cells collected at the indicated time points after their treatment with the protein synthesis inhibitor, emetine.

**Supplementary Figure 2.**
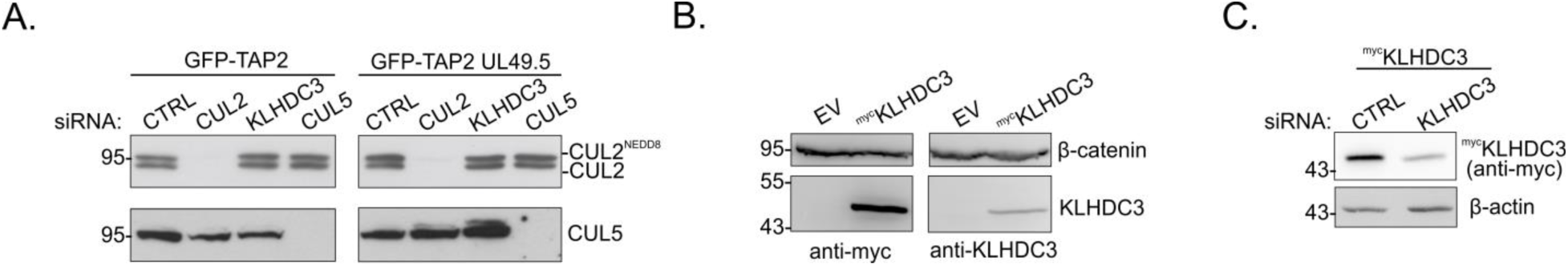
CRL2^KLHDC3^ complex is required for UL49.5-mediated TAP degradation. (A) MJS GFP-TAP2 cells stably expressing UL49.5 were transfected with the indicated siRNAs. The level of cullin 2 and cullin 5 was analyzed by immunoblotting. (B) MJS cells were transiently transfected with a plasmid encoding ^myc^KLHDC3 or an empty vector (EV). KLHDC3 was detected with anti-myc or anti-KLHDC3 antibodies. (C) MJS cells were transiently co-transfected with a plasmid encoding ^myc^KLHDC3 and siRNA targeting *KLHDC3*. The level of KLHDC3 was analyzed by immunoblotting with anti-myc antibodies.

**Supplementary Figure 3.**
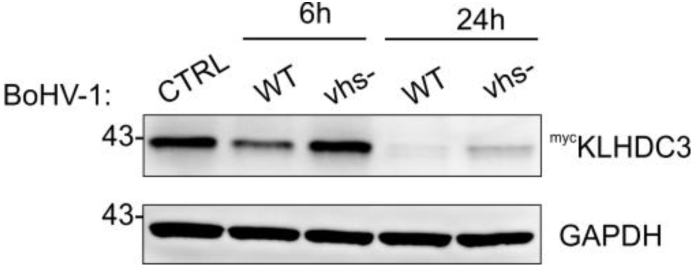
Interaction between UL49.5 and cullin 2 is time-regulated during BoHV-1 infection. MJS ^myc^KLHDC3 cells were infected with BoHV-1 WT or Δvhs (vhs-) mutant at an MOI of 10 for 6 or 24 hours. The level of KLHDC3 was analyzed by immunoblotting with anti-myc antibodies.

